# Assessing Confidence in Root Placement on Phylogenies: An Empirical Study Using Non-Reversible Models for Mammals

**DOI:** 10.1101/2020.07.31.230144

**Authors:** Suha Naser-Khdour, Bui Quang Minh, Robert Lanfear

## Abstract

Using time-reversible Markov models is a very common practice in phylogenetic analysis, because although we expect many of their assumptions to be violated by empirical data, they provide high computational efficiency. However, these models lack the ability to infer the root placement of the estimated phylogeny. In order to compensate for the inability of these models to root the tree, many researchers use external information such as using outgroup taxa or additional assumptions such as molecular-clocks. In this study, we investigate the utility of non-reversible models to root empirical phylogenies and introduce a new bootstrap measure, the *rootstrap*, which provides information on the statistical support for any given root position.

**Availability and implementation:** rootstrap support is implemented in IQ-TREE 2 and a tutorial is available at the iqtree webpage http://www.iqtree.org/doc/Rootstrap. In addition, a python script is available at https://github.com/suhanaser/Rootstrap. [phylogenetic inference, root estimation, bootstrap, non-reversible models]

## Main Text

The most widely used method for rooting trees in phylogenetics is the outgroup method. Although the use of an outgroup to root an unrooted phylogeny usually outperforms other rooting methods (Huelsenbeck, et al. 2002), the main challenge with this method is to find an appropriate outgroup (Watrous and Wheeler 1981; Maddison, et al. 1984; Smith 1994; Swofford, et al. 1996; Lyons-Weiler, et al. 1998; Milinkovitch and Lyons-Weiler 1998). Outgroups that are too distantly-related to the ingroup may have substantially different molecular evolution than the ingroup, which can compromise accuracy. And outgroups that are too closely related to the ingroup may not be valid outgroups at all.

It is possible to infer the root of a tree without an outgroup using molecular clocks (Huelsenbeck, et al. 2002; Drummond, et al. 2006). A strict molecular-clock assumes that the substitution rate is constant along all lineages, a problematic assumption especially when the ingroup taxa are distantly related such that their rates of molecular evolution may vary. Relaxed molecular-clocks are more robust to deviations from the clock-like behaviour (Drummond, et al. 2006), although previous studies have shown that they can perform poorly in estimating the root of a phylogeny when those deviations are considerable (Tria, et al. 2017).

Other rooting methods rely on the distribution of branch lengths, including Midpoint Rooting (MPR) (Farris 1972), Minimal Ancestor Deviation (MAD) (Tria, et al. 2017), and Minimum Variance Rooting (MVR) (Mai, et al. 2017). Such methods also assume a clock-like behaviour; however, they are less dependent on this assumption as the unrooted tree is estimated without it. Similar to inferring a root directly from molecular-clock methods, the accuracy of those rooting methods decreases with higher deviations from the molecular-clock assumption (Mai, et al. 2017).

Other less common rooting methods that can be used in the absence of outgroup are: rooting by gene duplication (Dayhoff and Schwartz 1980; Gogarten, et al. 1989; Iwabe, et al. 1989), indel-based rooting (Rivera and Lake 1992; Baldauf and Palmer 1993; Lake, et al. 2007), rooting the species tree from the distribution of unrooted gene trees (Allman, et al. 2011; Yu, et al. 2011), and probabilistic co-estimation of gene trees and species tree (Boussau, et al. 2013).

All the methods mentioned above, apart from the molecular-clock, infer the root position independently of the ML tree inference. The only existing approach to include root placement in the ML inference is the application of non-reversible models. Using non-reversible substitution models relaxes the fundamental assumption of time-reversibility that exists in the most widely used models in phylogenetic inference (Jukes and Cantor 1969; Kimura 1980; Hasegawa, et al. 1985; Tavaré 1986; Dayhoff 1987; Jones, et al. 1992; Tamura and Nei 1993; Whelan and Goldman 2001; Le and Gascuel 2008). This in itself is a potentially useful improvement in the fit between models of sequence evolution and empirical data. In addition, since non-reversible models naturally incorporate a notion of time, the position of the root on the tree is a parameter that is estimated as part of the ML tree inference. Since the incorporation of non-reversible models in efficient ML tree inference software is relatively new (Minh, et al. 2020), we still understand relatively little about the ability of non-reversible models to infer the root of a phylogenetic tree, although a recent simulation study has shown some encouraging results (Bettisworth and Stamatakis 2020).

Regardless of the rooting method and the underlying assumptions, it is crucial that we are able to estimate the statistical confidence we have in any particular placement of the root on a phylogeny. A number of previous studies have sensibly used ratio likelihood tests such as the Shimodaira-Hasegawa (SH) test (Shimodaira and Hasegawa 1999) and the Approximately Unbiased (AU) test (Shimodaira 2002) to compare a small set of potential root placements, rejecting some alternative root placements in favour of the ML root placement e.g.(Nardi, et al. 2003; Steenkamp, et al. 2006; Jansen, et al. 2007; Moore, et al. 2007; Williams, et al. 2010; Kocot, et al. 2011; Zhou, et al. 2011; Whelan, et al. 2015; Zhang, et al. 2018), these tests are still somewhat limited in that they do not provide the level of support the data have for a certain root position.

There is strong empirical evidence that molecular evolutionary processes are rarely reversible (Squartini and Arndt 2008; Naser-Khdour, et al. 2019), but few studies have explored the accuracy of non-reversible substitution models to root phylogenetic trees (Huelsenbeck, et al. 2002; Yap and Speed 2005; Williams, et al. 2015; Cherlin, et al. 2018; Bettisworth and Stamatakis 2020). Most studies that have looked at this question in the past have focused on either simulated datasets (Huelsenbeck, et al. 2002; Jayaswal, et al. 2011; Cherlin, et al. 2018) or relatively small empirical datasets (Yang and Roberts 1995; Yap and Speed 2005; Jayaswal, et al. 2011; Heaps, et al. 2014; Williams, et al. 2015; Cherlin, et al. 2018). In both cases, the addressed substitution models were nucleotide models, and to our knowledge, no study has yet investigated the potential of amino acid substitution models in inferring the root placement of phylogenetic trees.

In this paper, we focus on evaluating the utility of non-reversible amino acid and nucleotide substitution models to root the trees, and we introduce a new metric, the *rootstrap support value*, which estimates the extent to which the data support every possible branch as the placement of a root in a phylogenetic tree. Unlike previous studies that used Bayesian methods with non-reversible substitution models to infer rooted ML trees (Heaps, et al. 2014; Cherlin, et al. 2018), we will conduct our study in a Maximum likelihood framework using IQ-TREE (Minh, et al. 2020). A clear advantage of Maximum likelihood over the Bayesian analysis is that there is no need for a prior on the parameter distributions, which sometimes can affect tree inference (Huelsenbeck, et al. 2002; Cherlin, et al. 2018). Even though estimating the non-reversible model’s parameters by maximizing the likelihood function seems more computationally intensive than calculating posterior probabilities (Huelsenbeck, et al. 2002), the IQ-TREE algorithm is sufficiently fast to allow us to estimate root placements, with *rootstrap support* for very large datasets.

A recent study investigated the ability of non-reversible nucleotide models to infer the root placement of phylogenetic trees (Bettisworth and Stamatakis 2020). This study showed that IQ-TREE performs competitively with a new rooting tool, RootDigger. In most simulated datasets, IQ-TREE slightly outperformed RootDigger in terms of root placements, but no comparisons were made between RootDigger and IQ-TREE on empirical datasets. Although, RootDigger is significantly faster than IQ-TREE (Bettisworth and Stamatakis 2020), the former is limited to nucleotide substitution models. Since we are interested in both nucleotide and amino acid non-reversible models, we used IQ-TREE for tree and root inference in this study.

## Material and Methods

### The “Rootstrap” Support, and measurements of error in root placement

To compute rootstrap supports, we conduct a bootstrap analysis, i.e., resampling alignment sites with replacement, to obtain a number of bootstrap trees. We define the *rootstrap* support for each branch in the ML tree, as the proportion of bootstrap trees that have the root on that branch. Since the root can be on any branch in a rooted tree, the rootstrap support values are computed for all the branches including external branches. The sum of the rootstrap support values along the tree are always smaller than or equal to one. A sum that is smaller than one can occur when one or more bootstrap replicates are rooted on a branch that does not occur in the ML tree (Fig. 1).

**Figure 1.**
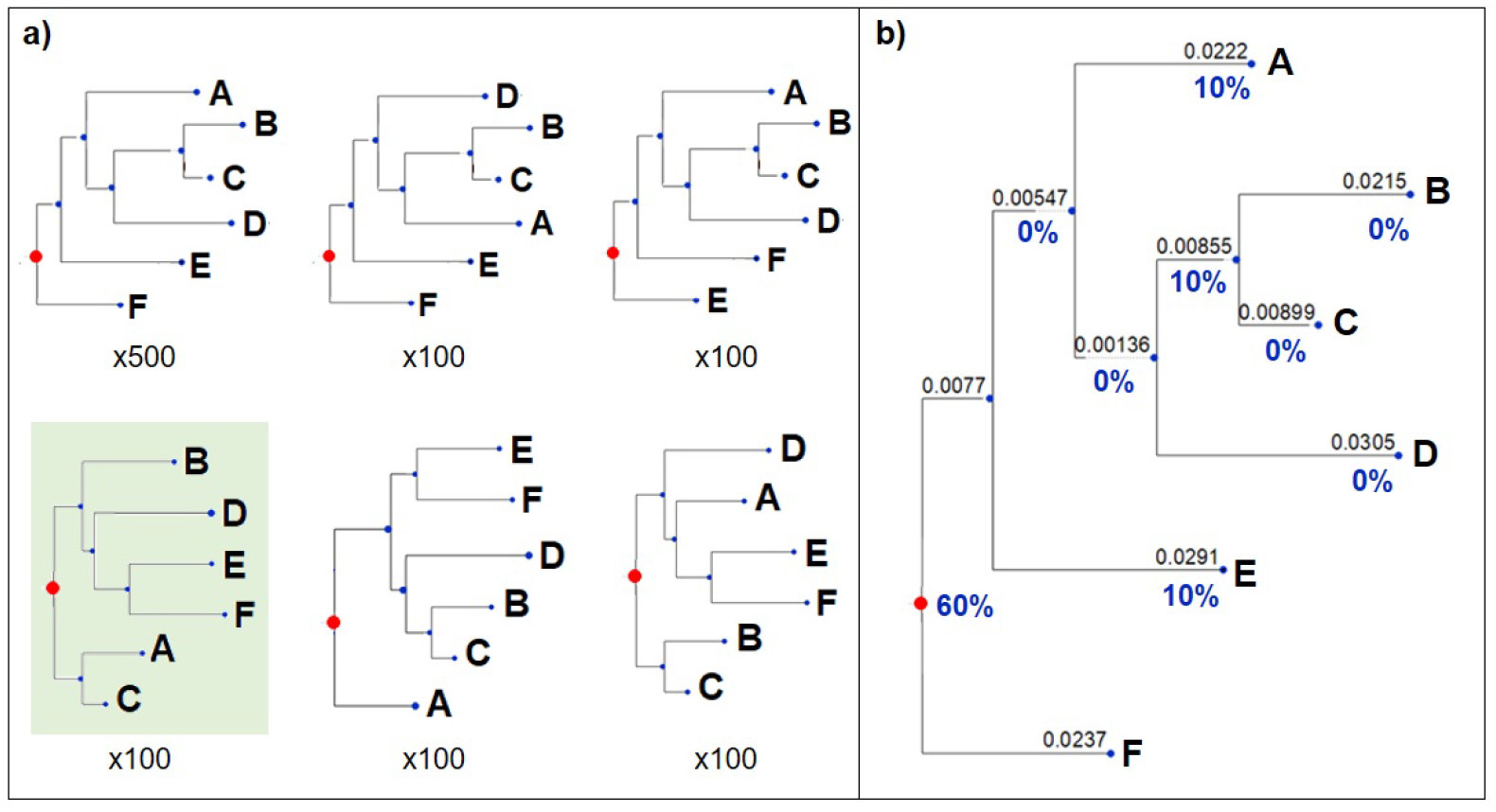
Illustration of the rootstrap concept. (a) The bootstrap replicates trees. (b) The ML tree with the rootstrap support values for each branch. Note that the sum of the rootstrap support values is less than 100% due to 100 bootstrap replicates trees (green) that have their root at a branch that does not exist in the ML tree.

By definition, the rootstrap support values for internal branches are bounded by the bootstrap support values at those branches. On the other hand, the rootstrap support values for tips (leaf branches) are bounded by 100%, as tips always appear in all the bootstrap trees.

If the true position of the root is known (e.g. in simulation studies) or assumed (e.g. in the empirical cases we present below), we can calculate additional measurements of the error of the root placement. We introduce two such measurements here: *root branchlength error distance* (rBED) and *root split error distance* (rSED). Since the non-reversible model infers the exact position of the root on a branch, we define the *root branchlength error distance* (rBED) as the range between the minimum and maximum distance between the inferred root position and the “true root” branch. If the true root is on the same branch as the ML tree root, then rBED will be between 0 and the distance between the ML tree root and the farthest point on that branch (Fig. 2). Since rBED is based on branch lengths only, it ignores the absolute number of splits between the ML tree root and the true root; and therefore, the rBED for the true root being on the same ML root branch can be bigger than the rBED for the true root being on a different branch (e.g. Fig. 2). In order to account for the number of splits (nodes) between the ML tree root and the true root, we define *root split error distance* (rSED) as the number of splits between the ML root branch and the branch that is believed to contain the true root (Fig. 2).

**Figure 2.**
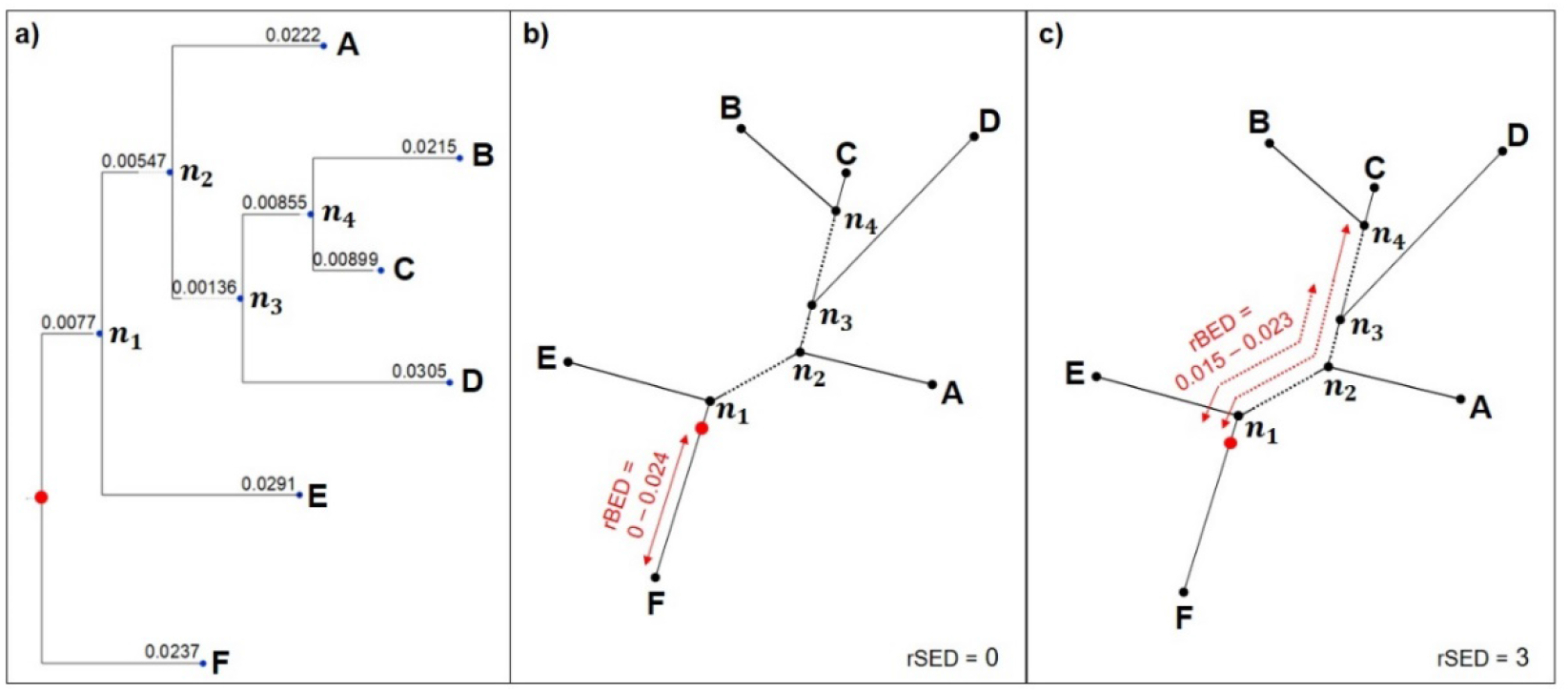
An example to illustrate the root error distance. (a) the ML rooted tree, (b) the root branch-length error distance (rBED) if the true root is believed to be on the same ML root branch (rSED = 0), (c) the rBED if the true root is believed to be on the branch between D and the clade of C + B (rSED = 3).

The rootstrap, rBED, and rSED assess different aspects of the root placement. While the rootstrap offers an indication of the support that the data have for a certain branch to be the root branch, rBED and rSED provide an estimation to the accuracy of the method in estimating the exact root position if the root position is known or assumed in advance. In other words, the rootstrap value is a measure for the robustness of the root placement given the model and the data and can be used on any dataset regardless of whether the true root position is known, while rBED and rSED are measures of the accuracy of the non-reversible model to find the root placement given the data, and require the root position to be known or assumed in advance.

### Empirical Datasets

Because non-reversible amino acid models require the estimation of a large number of parameters, and because we suspected that the information in any such analysis on the placement of the root branch of a tree might be rather limited, we searched for empirical datasets that met a number of stringent criteria:

1. Existence of both DNA and amino acid multiple sequences alignments (MSA) for the same loci.
2. Genome-scale MSAs to ensure that the MSAs have as much information as possible with which to estimate the non-reversible models’ free parameters and the root position. Since we do not know the number of sites required to correctly infer the rooted ML tree, we define 100,000 sites as the minimum number of required sites. This also allows us to subsample the dataset to explore the ability of smaller datasets to infer root positions.
3. Highly-curated alignments: since the quality of the inferred phylogeny is highly dependent on the quality of the MSA (Philippe, et al. 2011), we focussed on datasets that were highly-curated for misalignment, contamination, and paralogy.
4. Existence of several clades for which there is a very strong consensus regarding their root placement. Since we are interested in evaluating the performance of non-reversible models to infer root placements in an empirical rather than a simulation context, we need to identify monophyletic sub-clades for which we can be almost certain about their root position. This enables us to divide the dataset into non-overlapping sub-clades for which we are willing to assume we know the root positions. Furthermore, we define the minimum number of taxa in each sub-dataset as five.

We initially identified a number of genome-scale datasets that contained large numbers of nucleotide and amino acid MSAs. In many cases, it was difficult to determine whether these alignments had been rigorously curated, and even more challenging to find datasets for which the root position of a number of subclades could be assumed with confidence. The only dataset that met all of our criteria was a dataset of placental mammals with 78 ingroup taxa and 3,050,199 amino acids (Wu, et al. 2019). This dataset was originally published as an MSA (Liu, et al. 2017) based on very high-quality sequences from Ensembl, NCBI, and GenBank databases. After receiving detailed critiques for potential alignment errors (Gatesy and Springer 2017), the dataset was further processed to remove potential sources of bias and error, and an updated version of the dataset was recently published (Wu, et al. 2018). The fact that this alignment comes from one of the most well-studied clades on the planet, has been independently curated and critiqued by multiple groups of researchers and includes truly genome-scale data, makes it ideally suited for our study. The curated alignments can be found on figshare (https://figshare.com/s/622e9e0a156e5233944b) under the name “Wu_2018_aa” and “Wu_2018_dna” for the amino-acid and nucleotide alignments respectively.

### Selecting Clades with a Well-Defined Root

Since our main objective in this study is to evaluate the effectiveness of non-reversible models and the rootstrap value in estimating and measuring the support for a given root placement on empirical datasets, we must identify a collection of sub-clades of the larger mammal dataset for which it is reasonable to assume a root position. We acknowledge, of course, that outside a simulation framework it is not possible to be certain of the position of the root position of a clade. Nevertheless, it is possible to identify clades for which the position of the root is well supported and non-controversial, thus minimising the chances that the assumption of a particular root position is incorrect. To achieve this, we analysed the root position of each order and superorder in the dataset, and defined “*well-defined clades*” that fulfilled **all** of the following criteria:

1. It contains at least five taxa. This ensures that the probability of obtaining a random ML rooted tree to be at most 0.95%. For clades with four taxa, there are 15 different rooted topologies, and therefore a 6.7% probability to get any particular root position by chance. On the other hand, for clades with at least five taxa, there are at least 105 different rooted topologies and maximum probability of 0.95% to randomly get a particular root position by chance.
2. The bootstrap support for the branch leading to that clade in the phylogenetic tree calculated from the whole dataset is 100%: since the bootstrap value indicates the support the data have for a certain branch, we also require 100% support for the first direct descendants in the clade (Appendix Fig. A.1). This requirement ensures that there is strong support in the dataset for the root position of the clade when the entire dataset is rooted with an outgroup.
3. The site concordance factor (sCF) for the first direct descendants in the clade is significantly greater than 33%. The site Concordance Factor (sCF) is calculated by comparing the support of each site in the alignment for the different arrangements of quartet around a certain branch. In other words, an sCF of 33% means equal support for any of the possible arrangements. Therefore, we require that the sCF of the deepest two levels of branches leading to that clade is significantly greater than 33%. Moreover, we require that the gene Concordance Factor (gCF) for the first direct descendants in the clade to be significantly greater than 33% of the sum of the gene concordance factor and the two Discordance Factors (gDF1 and gDF2). The gCF of a branch is calculated as the proportion of gene trees containing that branch, and gDFs are calculated as the proportion of gene trees containing one of the two other resolutions of that branch. Since for each branch in a bifurcating tree there are three possible arrangements of clades around that branch, we ignore all gene trees that do not contain one of these arrangements (e.g. gene trees that contribute to neither the gCF nor the gDFs). Although there is no threshold regarding the required proportion of genes concordant with a certain branch, for convenience, we define branches with gCF significantly greater than 33% of the sum gCF+gDF1+gCF2 as branches that are concordant with enough genes in the alignment (Minh, et al. 2020). To test whether the sCF and the gCF are significantly greater than 33%, we use a simple binomial test with a success probability of 0.33. The gCF,gDF1,gCF2 and sCF values are based on the tree estimated from the amino acid dataset.
4. At least 95% of the studies that have been published in the last decade support this clade: we searched google scholar for all published papers since 2009 that determine the root of the addressed clade. We then checked if at least 95% of those papers agree that the root position of the clade matches that in the ML tree we estimate from the whole dataset (see supplementary material).

### Estimating unrooted Phylogenies

For the whole nucleotide and amino-acid datasets with ingroup and outgroup taxa, we inferred the unrooted phylogeny using IQ-TREE2 (Minh, et al. 2020) with the best-fit fully partitioned model (Chernomor, et al. 2016) and edge-linked substitution rates (Duchene, et al. 2020). We then determined the best-fit reversible model for each partition using ModelFinder (Kalyaanamoorthy, et al. 2017). See the algorithm for finding well-defined clades in Appendix Algorithm A.1.

### Estimating Rooted phylogenies

For each well-defined clade, we first removed all other taxa from the tree and then sought to infer the root of the well-defined clade using non-reversible models without outgroups. Using the best partitioning scheme from the reversible analysis, we inferred the rooted tree for each well-defined clade with the non-reversible models for amino acid (NR-AA) and nucleotide (NR-DNA) sequences (Minh, et al. 2020). This approach fits a 12-parameter non-reversible model for DNA sequences, and a 380-parameter non-reversible model for amino acids. Details of the command lines used are provided in the supplementary material section “Algorithm A.2”. Each analysis returns a rooted tree. We performed 1000 non-parametric bootstraps of every analysis to measure the rootstrap support.

To assess the performance of the rootstrap and the ability of non-reversible models to estimate the root of the trees on smaller datasets, we also repeated every analysis on subsamples of the complete dataset. For each well-defined clade, we performed analysis on the complete dataset (100%) as well as datasets with 10%, 1% and 0.1% of randomly-selected loci from the original alignment.

### The confidence set of root branches using the Approximately Unbiased test

In addition to the rootstrap support, we calculate the confidence set of all the branches that may contain the root of the ML tree using the Approximately Unbiased (AU) test (Shimodaira 2002). To do this, we re-root the ML tree with all possible placements of the root (one placement for each branch) and calculate the likelihood of each tree. Using the AU test, we then ask which root placements can be rejected in favour of the ML root, using an alpha value of 5%. We define the *root branches confidence set* as the set of root branches that are not rejected in favour of the ML root placement. An important difference between the AU test and the rootstrap support is that the AU test is conditioned on a single ML tree topology, but the rootstrap support is not. Because of this, they provide quite different information about the position of the root. The AU test assumes that the ML tree topology is true, and then seeks to determine the confidence set of root placements conditioned on that topology. The confidence set for the AU test will always therefore contain at least the ML root branch. The rootstrap does not assume any particular topology, and instead asks how many times a particular root position appears across a set of bootstrap replicates. Because of this, it is possible for every branch in the ML topology to receive 0% rootstrap support. This can occur if none of the branches in the ML topology appear as the root branch in any of the bootstrap topologies.

### Reducing systematic bias by removing third codon positions and loci that fail the MaxSym test

As it is common in many phylogenetic analyses to remove third codon positions from the alignment (Swofford, et al. 1996), we wanted to assess the effect of removing third codon positions on the root inference and the rootstrap values in nucleotide datasets. For that purpose, we remove all the third codon positions from the nucleotide alignments and re-ran the analysis using the NR-DNA model.

Moreover, although the NR-AA and NR-DNA models relax the reversibility assumption, they still assume stationarity and homogeneity. To reduce the systematic bias produced by violating these assumptions, we used the MaxSym test (Naser-Khdour, et al. 2019) to remove loci that violate those assumptions in the nucleotide and amino acid datasets, and then re-ran all analyses as above.

### Applying the methods to two clades whose root position is uncertain

In addition to the well-defined clades, we used the methods we propose here to infer the root of two clades of mammals whose root position is controversial; Chiroptera and the Cetartiodactyla.

There is a controversy around the root of the Chiroptera (bats) in literature. The two most popular hypotheses are: 1) the Microchiroptera-Megachiroptera hypothesis; where the root is placed between the Megachiroptera, which contains the family Pteropodidae, and the Microchiroptera, which contains all the remaining Chiroptera families. This hypothesis is well supported in the literature (Agnarsson, et al. 2011; Meredith, et al. 2011). However, more recent studies seem to provide less support for this hypothesis; 2) the Yinpterochiroptera-Yangochiroptera hypothesis, in which the Yangochiroptera clade includes most of Microchiroptera and the Yinpterochiroptera clade includes the rest of Microchiroptera and all of Megachiroptera. There is growing support for this hypothesis in the literature (Meganathan, et al. 2012; Tsagkogeorga, et al. 2013; Ren, et al. 2018; Reyes-Amaya and Flores 2019).

Similar to Chiroptera, the root of Cetartiodactyla remains contentious in the literature. The three main hypotheses regarding the root of Cetartiodactyla are: 1) Tylopoda as the sister group for all other cetartiodactylans; 2) Suina as the sister group for all other cetartiodactylans; 3) the monophyletic clade containing Tylopoda and Suina as the sister group for all other cetartiodactylans.

To ascertain whether certain sites or loci had very strong effects on the placement of the root we follow the approach of Shen et. al. (Shen, et al. 2017) and calculate the difference in site-wise log-likelihood scores (ΔSLS) and gene-wise log-likelihood scores (ΔGLS) between the supported root positions for each clade. Moreover, we analysed subsamples of each dataset to test the limits of using non-reversible models to root trees with smaller datasets.

## Results

### Inference of the mammal tree and selection of well-defined clades

The trees inferred from the whole datasets with the nucleotide-reversible model and the amino-acid-reversible model (Appendix Fig. A.2, Appendix Fig. A.3, Appendix Table A.2) are consistent with the published tree (Liu, et al. 2017). Five clades met all the criteria of well-defined clades, namely, Afrotheria, Bovidae, Carnivora, Myomorpha, and Primates in both amino acid and nucleotide datasets (see Appendix Table A.1 and Appendix Table A.2). Trees in Newick format can be found on github: https://github.com/suhanaser/Rootstrap/tree/master/trees

### High accuracy of the AA non-reversible model in inferring the root

Using NR-AA, we inferred the correct root with very high rootstrap support for all five well-defined clades when all loci were used (Appendix Table A.3). Moreover, for all the five clades, the true root was the only root placement in the confidence set of the AU test. The average running time of the NR-AA model (model estimation + tree search + bootstrap + root inference) is 929 hrs on one core 2.6GHz CPU. However, using the optimal number of cores for each dataset reduced the average running time to 43.5 hrs per dataset.

Our results show that using only 10% of the sites in the amino acid alignments (around 300,000 alignment columns) still gave very high rootstrap support values (> 98%) for four of the five well-defined clades (Fig. 3) with no correlation between rSED and rBED and the size of the dataset (Table A.3). Moreover, in three of five well-defined clades, 1% of the sites (around 30,000 alignment columns) was enough to give a very high rootstrap support value for the assumed correct root placement. Using only 0.1% of the sites (around 3000 alignment columns) decreased the rootstrap support value noticeably in all datasets (Appendix Table A.3). These values are shown for each dataset in Figure 3, where the X-axis is plotted in terms of parsimony-informative sites to allow for a more direct comparison between datasets, and to assist those applying these methods in deciding whether to use them on their own data. Although the rootstrap support for the true root improves as the number of parsimony-informative sites increase, in some datasets (e.g. Afrotheria nucleotide dataset) this is not the case (Fig. 3).

**Figure 3.**
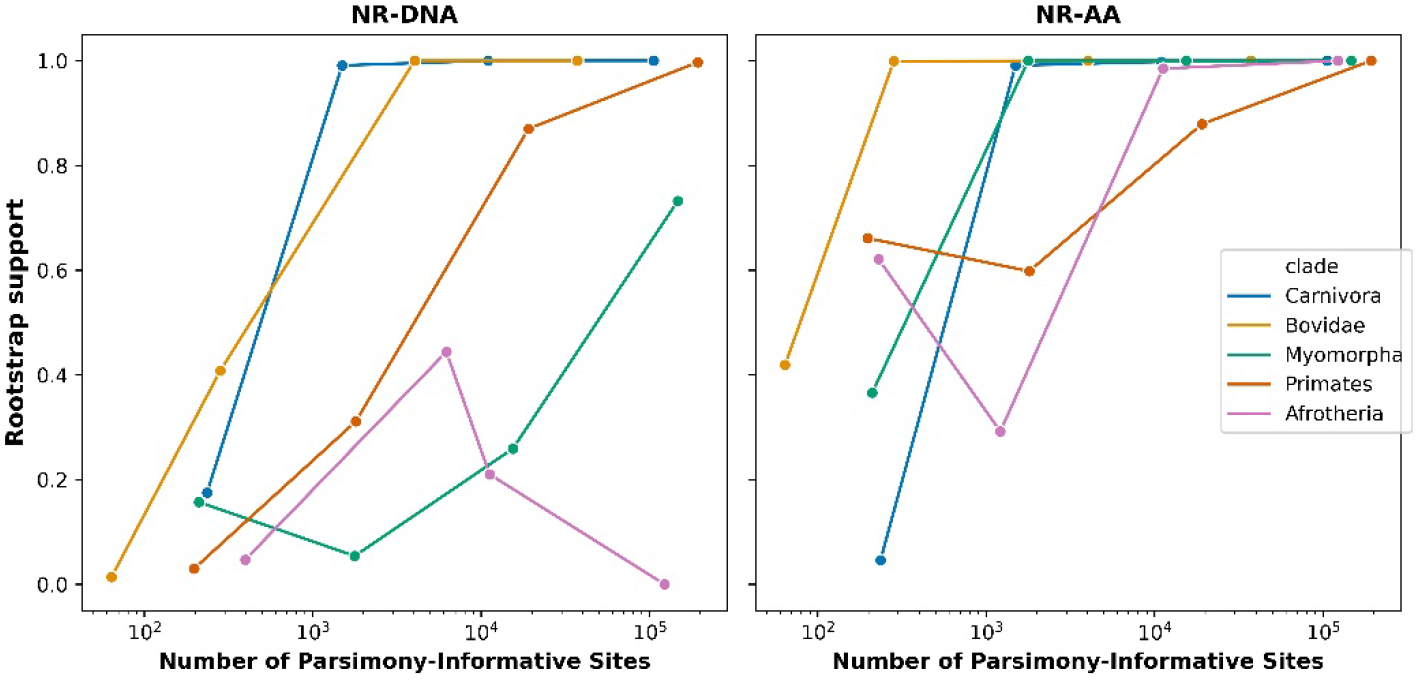
The rootstrap support value for each clade as a function of the number of parsimony-informative sites.

The non-reversible amino acid models were strongly preferred to the reversible models on the complete datasets (BIC values were 93943 to 235958 units better for the non-reversible models), and for the datasets with 10% of loci subsampled (BIC values were 3577 to 15082 units better for the non-reversible models), but the opposite was true for the datasets 1% and 0.1% of the loci subsampled (e.g. BIC values were between 2102 and 2712 units worse for the non-reversible models for the 0.1% subsampled datasets; see Table A.7 for full results).

### Poor performance of the DNA non-reversible model in inferring the root

We correctly inferred the root for four out of the five nucleotide datasets with the NR-DNA model, when all loci were used. However, the rootstrap support was generally lower than in the amino-acid datasets (Fig. 3, Appendix Tables A.3 and A.4). Similar to amino-acid datasets, there is no correlation between rSED and rBED and the size of the dataset (Table A.4). The average running time of the NR-DNA model (model estimation + tree search + bootstrap + root inference) is 35.7 hrs on one core 2.6GHz CPU and 4 hours when the optimal number of cores for each dataset were used.

In contrast to the NR-AA model, there is no conclusive preference for the NR-DNA model over the reversible DNA model for the datasets we analysed (Table A.8). In fact, The BIC values of the NR-DNA models are always worse than reversible models regardless to the size of the nucleotide dataset except for three clades when all loci were included (Table A.8). In two of the datasets (Myomorpha and Primates) were the NR-DNA model was better than the reversible model the root placement was inferred correctly with high rootstrap support (>95%). In fact, the Afrotheria nucleotide dataset is the only dataset in which the non-reversible model was better than the reversible model but the root placement was inferred incorrectly.

Our results show that removing the third codon positions does not improve the rootstrap support value. In contrast, in some datasets removing third codon positions decreased the rootstrap support value and increased the rSED (Table 1).

**Table 1.** Rootstrap support and rSED values in whole nucleotide datasets and nucleotide datasets without third codon positions.

| Clades | All loci |  | Without 3rd |  |
| --- | --- | --- | --- | --- |
|  | rootstrap | rSED | rootstrap | rSED |
| Afrotheria | 0.0% | 2 | 0.0% | 2 |
| Primates | 99.7% | 0 | 90.1% | 0 |
| Myomorpha | 73.2% | 0 | 15.8% | 1 |
| Carnivora | 100.0% | 0 | 100.0% | 0 |
| Bovidae | 100.0% | 0 | 82.5% | 0 |

### Removing loci that violate the stationarity and homogeneity assumptions improves the rootstrap support

As expected, our results show that removing loci that fail the MaxSym test improves the rootstrap support values when the rootstrap support value was less than 100% and/or the root placement was inferred incorrectly, as the case in some nucleotide datasets (Table 2).

**Table 2.** Rootstrap support values in whole datasets and datasets with loci that passed the MaxSym test only.

| Clade | Amino Acid |  | Nucleotide |  |
| --- | --- | --- | --- | --- |
|  | all loci | Passed MaxSym | all loci | Passed MaxSym |
| Afrotheria | 100.0% | 100.0% | 0.0% | 8.4% |
| Primates | 100.0% | 100.0% | 99.7% | 99.9% |
| Myomorpha | 100.0% | 100.0% | 73.2% | 88.3% |
| Carnivora | 100.0% | 100.0% | 100.0% | 100.0% |
| Bovidae | 100.0% | 100.0% | 100.0% | 100.0% |

### Microchiroptera-Megachiroptera or Yinpterochiroptera-Yangochiroptera?

Using the whole amino acid dataset, our results show 65.5% rootstrap support for the Yinpterochiroptera-Yangochiroptera hypothesis and 23.2% for the Microchiroptera - Megachiroptera hypothesis. The remaining11.3% of the rootstrap support goes to supporting the branch leading to Rhinolophoidea as root branch of the bats (Fig. 4). Removing amino acid loci that fail the MaxSym test (110 loci) gives similar results, with 65.9% rootstrap support for the Yinptero-Yango hypothesis and 25.6% rootstrap support for the Micro-Mega hypothesis. In both cases, the AU test could not reject any of the three root positions that received non-zero rootstrap support (Appendix Table A.5).

**Figure 4.**
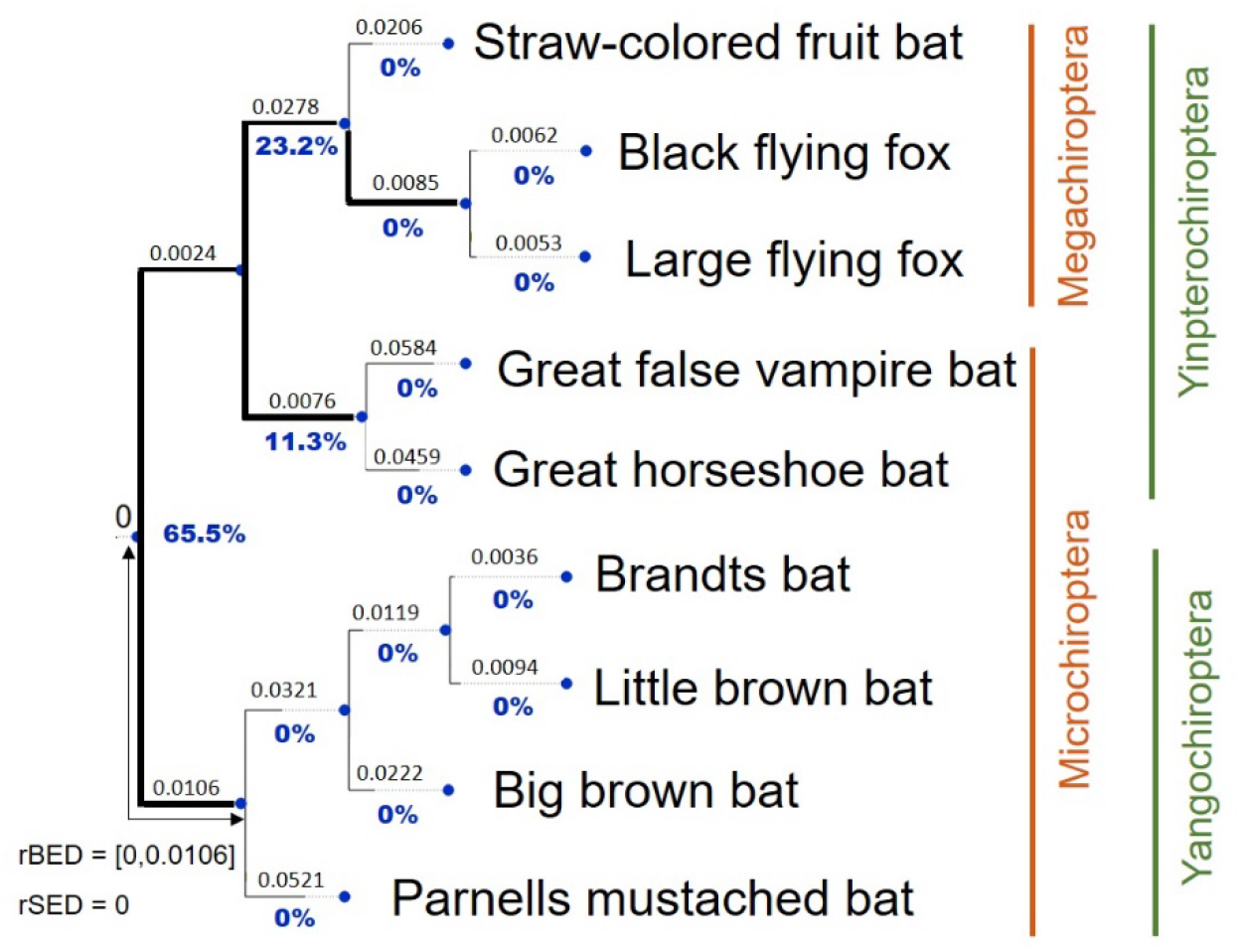
The ML rooted tree as inferred from the whole Chiroptera amino acid dataset. Bold branches are branches in the AU confidence set. Blue values under each branch are the rootstrap support values.

Using the NR-DNA model gives 100% rootstrap support for the Yinptero-Yango hypothesis, and we can confidently reject the Micro-Mega hypothesis in favour of the Yinptero-Yango hypothesis using the AU test (Appendix Fig. A.4). Yet, removing nucleotide loci that fail the MaxSym test (∼25% of the loci) decreases the support for the Yinptero-Yango hypothesis to 90.1%, although we can still confidently reject the Micro-Mega hypothesis using the AU test (Appendix Table A.5).

Interestingly, when we randomly subsample 10%, 1%, and 0.1% of the loci in the nucleotide dataset, we consistently get the Yinptero-Yango hypothesis as the ML tree and the solely rooted topology in the AU confidence set (Appendix Table A.5). Moreover, the rootstrap support value for the Yinptero-Yango hypothesis increases and the rootstrap support value for the Micro-Mega hypothesis decreases as more parsimony-informative sites are added to the alignment, for both nucleotide and amino acid datasets (Fig. 5, Appendix Table A.5). These results are consistent with previous studies that used smaller datasets (Appendix Figure A.10)

**Figure 5.**
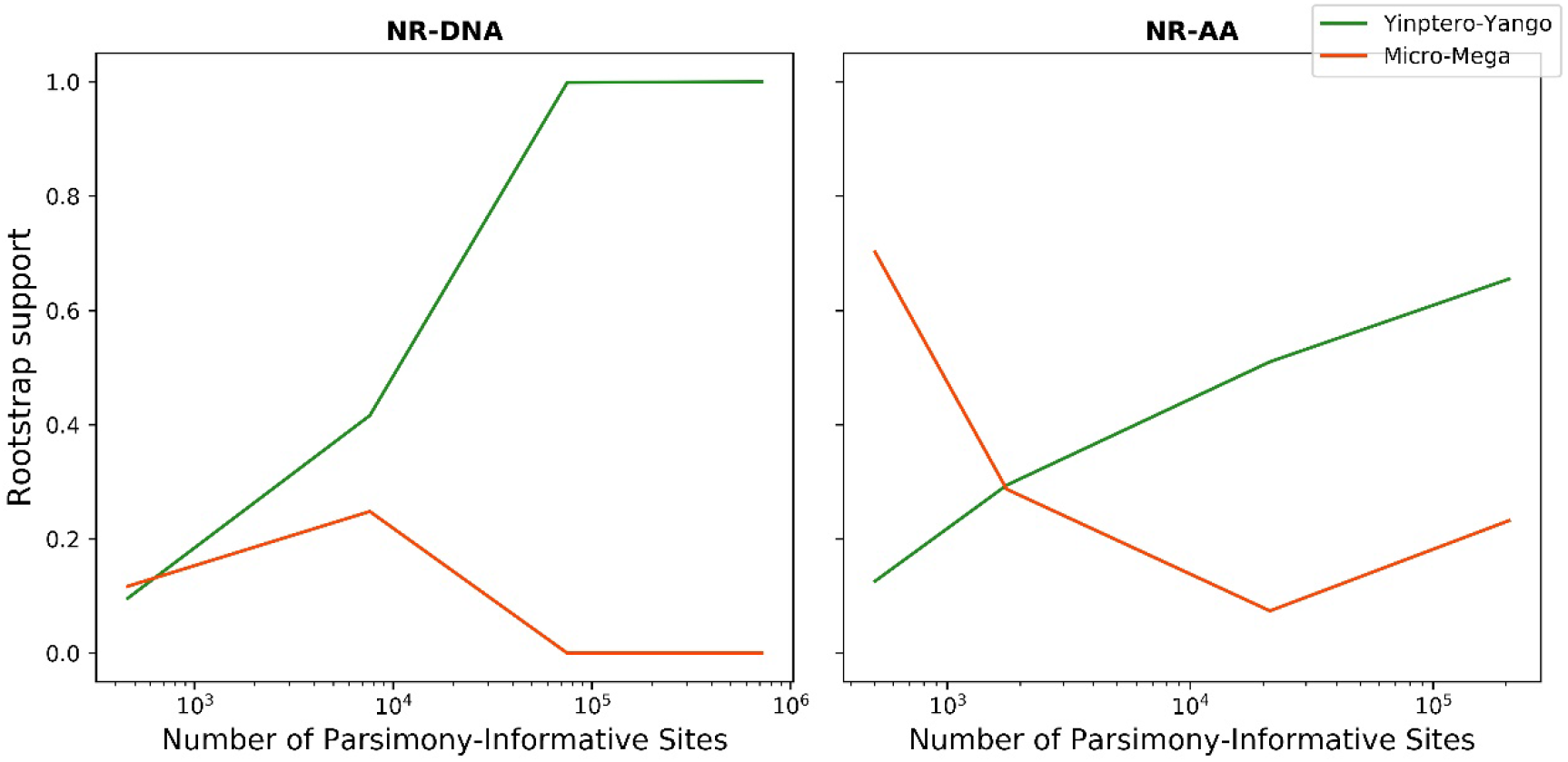
Rootstrap support value as a function of the number of parsimony-informative characters in the Chiroptera nucleotide and amino acid datasets using the NR-DNA model (to the left) and the NR-AA model (to the right).

The ΔGLS and ΔSLS values (Shen, et al. 2017) reveal that approximately half of the nucleotide and amino acid loci prefer the Yinptero-Yango hypothesis while the other half prefers Micro-Mega hypothesis. Furthermore, slightly less than half of the nucleotide sites prefer the Yinptero-Yango hypothesis. However, more than two-thirds of the amino acid sites prefer the Yinptero-Yango hypothesis (Appendix Fig. A.5). The distributions of ΔGLS and ΔSLS (Appendix Fig. A.6) show that a small proportion of the amino acid loci (∼1%) have very strong support for the Micro-Mega hypothesis, and removing those loci from the alignment increased the rootstrap support for the Yinptero-Yango hypothesis to 76.6%. Nonetheless, both root placements are still in the confidence set of the AU test (Appendix Table A.5) with the amino acid dataset. On the other hand, removing nucleotide loci with the highest absolute ΔGLS value still gives the Yinptero-Yango hypothesis as the ML tree and the sole topology in the AU confidence set. Although the nucleotide data show a clear preference to the Yinptero-Yango hypothesis, in terms of BIC scores, the NR-DNA model preforms worse than reversible models in all datasets except for the dataset where we removed loci that failed the MaxSym test (Table A.5). On the other hand, the NR-AA performs better than reversible models in big datasets (Table A.5). Yet, the amino acid data do not allow us to distinguish between the two leading hypotheses for the placement of the root of the Chiroptera based on rooting with non-reversible models (Table A.5).

### The ambiguous root of Cetartiodactyla

The ML tree inferred with the whole amino acid dataset places the clade containing Tylopoda (represented by its only extant family; Camelidae) and Suina as the sister group to all other cetartiodactylans with 71.8% rootstrap support (Fig. 6). Yet, The AU test did not reject Tylopoda alone as the sister group to all other cetartiodactylans. On the other hand, the ML tree inferred with the whole nucleotide dataset places Tylopoda as the only sister group to all other cetartiodactylans with 71.0% rootstrap support, and we can confidently reject the Tylopoda + Suina hypothesis using the AU test (Appendix Fig. A.7).

**Figure 6.**
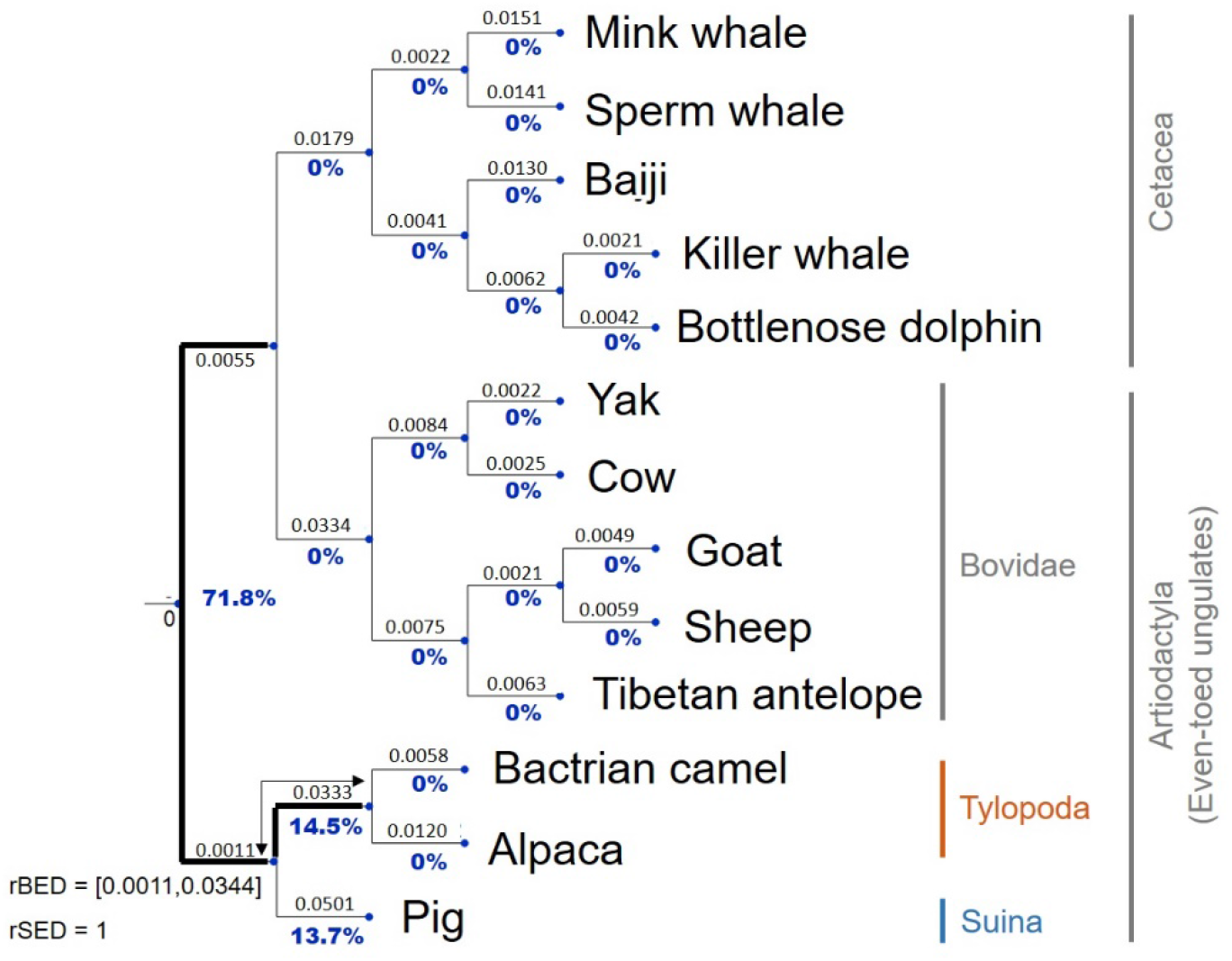
The ML rooted tree of as inferred from the whole Cetartiodactyla amino acid dataset. Bold branches are branches in the AU confidence set. Blue values under each branch are the rootstrap support values.

Removing the amino acid loci that failed the MaxSym test (∼1%) still places Tylopoda + Suina as the sister group to all other cetartiodactylans, yet, it decreases the rootstrap support for the Tylopoda + Suina hypothesis to 63.3% and increases the rootstrap support for the Tylopoda hypothesis to 28.5%. However, we still cannot reject either of the hypotheses using the AU test (Appendix Table A.6).

Removing the nucleotide loci that failed the MaxSym test (∼1%) still places Tylopoda as the only sister group to all other cetartiodactylans and the only rooted topology in the AU confidence set. However, it decreases the rootstrap support for the Tylopoda hypothesis to 68.7% and increases the rootstrap support for the Tylopoda + Suina hypothesis to 20.1% (Appendix Table A.6).

The results from the subsample datasets are mixed (Fig. 7). Analyses on smaller datasets show no clear pattern in the placement of the root (Appendix Table A.6), leading us to conclude only that the analyses of the whole dataset is likely to provide the most accurate result, but that it is plausible that adding more data may lead to different conclusions in the future.

**Figure 7.**
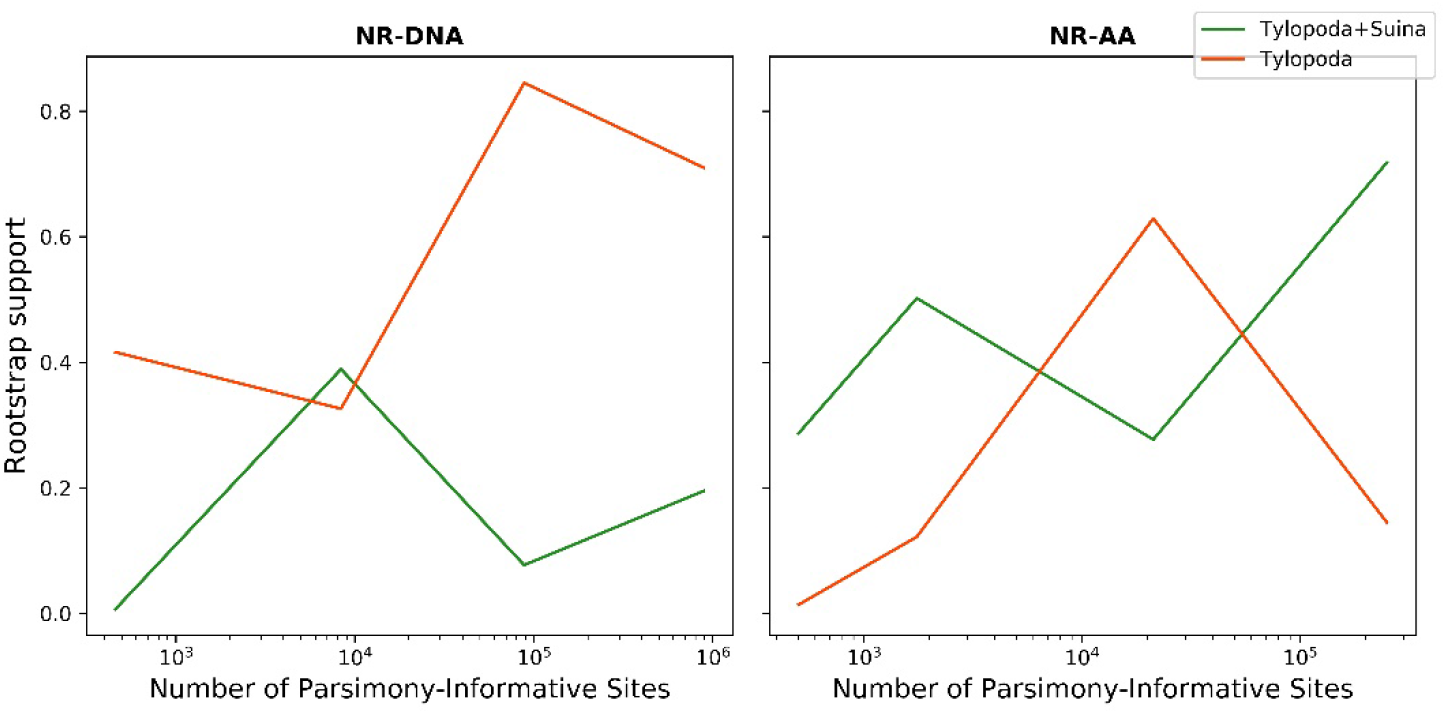
rootstrap support value as a function of the number of parsimony-informative characters in the Cetartiodactyla nucleotide and amino acid datasets using the NR-DNA model (to the left) and the NR-AA model (to the right).

ΔGLS analyses reveal that approximately, half of the amino acid and nucleotide loci favour the Tylopoda+Suina hypothesis, while the other half of loci favour the Tylopoda hypothesis (Appendix Figs. A.8-9). On the other hand, two-thirds of the amino acid sites and more than 80% of the nucleotide sites favour the Tylopoda+Suina hypothesis. Removing 1% of the amino acid loci with the highest absolute ΔGLS values still places Tylopoda + Suina as the sister group to all other cetartiodactylans. However, the rootstrap support of the Tylopoda + Suina decreased to 63.2% and the rootstrap support for the Tylopoda hypothesis remains approximately the same (∼14.5%), while the rootstrap support for the Suina hypothesis increases from 13.7% to 22.4%. Yet, both the Tylopoda + Suina hypothesis and the Tylopoda hypothesis are in the confidence set of the AU test, while the Suina hypothesis is rejected by the AU test (Appendix Table A.6).

Removing 1% of the nucleotide loci with the highest absolute ΔGLS values gives the Tylopoda+Suina as the sister group to all other cetartiodactylans with 39.7% rootstrap support. However, the solely rooted topology in the AU confidence set is the topology in which the root is placed on the branch leading to Suina (Appendix Table A.6). Similar to Chiroptera and the well-defined clades, NR-AA model preforms, in terms of the BIC score, better than reversible models in big amino-acid datasets, while the NR-DNA performs worse than reversible models in all datasets (Table A.6). We conclude that neither the nucleotide nor the amino acid data are adequate to infer the root placement of Cetartiodactyla with non-reversible models.

## Discussion

In this paper, we introduced a new measure of support for the placement of the root in a phylogenetic tree, the rootstrap support value, and applied it to empirical amino acid and nucleotide datasets inferred using non-reversible models implemented in IQ-TREE (Minh, et al. 2020). The rootstrap is a useful measure because it can be used to assess the statistical support for the placement of the root in any rooted tree, regardless of the rooting method. In a Maximum Likelihood setting, interpretation of the rootstrap support is similar to the interpretation of the classic nonparametric bootstrap. In a Bayesian setting, the same procedure could be used to calculate the posterior probability of the root placement given a posterior distribution of trees. It is noteworthy that the rootstrap support value is not a measure of the accuracy of the root placement and therefore should not be interpreted as such. However, it provides information about the robustness of the root inference with regard to resampling the data. This interpretation is consistent with the interpretation of the nonparametric bootstrap (Holmes 2003) but with regard to the root placement instead of the whole tree topology.

In addition to the rootstrap support value, we introduced another two metrics; the root branch-length error distance (rBED), and the root split error distance rSED. Similar to the rootstrap metric, these additional metrics can be used in with any approach that generates rooted phylogenetic trees. We note that both metrics require the true position of the root to be known (or assumed) and that the rBED requires the rooting method to be able to accurately place the root in a specific position of the root branch.

In this study, we used these and other methods to assess the utility of non-reversible models to root phylogenetic trees in a Maximum Likelihood framework. We focussed on applying these methods to a large and very well curated phylogenomic dataset of mammals, as the mammal phylogeny provides perhaps the best opportunity to find clades for which the root position is known with some confidence. As expected, our results show an exponential increase in the rootstrap support for the true root as we add more information to the MSA. Our results suggest that non-reversible amino-acid models are more useful for inferring root positions than non-reversible DNA models. One explanation for this difference between the NR-DNA and the NR-AA models is the bigger character-state space of the NR-AA models. These models have 400 parameters (380 rate parameters and 20 amino acid frequencies) whereas NR-DNA models have only 16 parameters (12 rate parameters and 4 nucleotide frequencies). This could allow the NR-AA model to capture the evolutionary process better than the NR-DNA model, potentially providing more information on the root position of the phylogeny. This hypothesis requires some further exploration though, and we note that the actual character-space of amino acids is much smaller than accommodated in NR-DNA models due to functional constraints on protein structure (Dayhoff, et al. 1978).

Another explanation for the difference in performance between the NR-AA and NR-DNA models is that higher compositional heterogeneity in nucleotide datasets may bias tree inference. The fact that each amino acid can be specified by more than one codon, and that synonymous substitutions are more frequent than non-synonymous substitutions, makes amino acid datasets less compositionally heterogeneous than nucleotide datasets. In principle, this bias can be alleviated by removing loci that violate the stationarity and homogeneity assumptions (Naser-Khdour, et al. 2019). Our results suggest that this may be the case for the datasets we analysed: we show that removing loci that violate the stationarity and homogeneity assumptions improves the accuracy and statistical support for the placement of the root. This is not surprising since the robustness of the rootstrap, similar to the bootstrap, relies on the consistency of the inference method, so removing systematic bias should improve its performance.

We used the non-reversible approach to rooting trees along with the rootstrap support to assess the evidence for different root placements in the Chiroptera and Cetartiodactyla. Using the amino acid datasets we found that in both cases, although there tended to be higher rootstrap support for one hypothesis, neither of the current hypotheses for either dataset could be rejected. These results emphasize the importance of the rootstrap support value as a measure of the robustness of the root estimate given the data. In both the Chiroptera and Cetartiodactyla datasets the root placement varied among subsamples of the dataset, and the rootstrap support reflects this uncertainty. However, checking the stability of root placement estimate by randomly subsampling from the whole Chiroptera dataset show an obvious trend towards the Yinpterochiroptera-Yangochiroptera hypothesis as the dataset increases in size. This trend is consistent with a small number of influential sites or loci having their signal progressively drowned out in favour of the Yinpterochiroptera-Yangochiroptera hypothesis as more data are added to the alignment. In both the Chiroptera and Cetartiodactyla cases, the amino acid data is inadequate to distinguish between certain root placements. On the other hand, in both the Chiroptera and Cetartiodactyla, the nucleotide datasets appear to show stronger support for a single root placement.

Comparing BIC scores of reversible and non-reversible models show that in most of the nucleotide datasets the reversible model was a much better fit to the data than the NR-DNA model. This is likely due to the limitations of the method we used to infer the NR-DNA model. Specifically, when inferring the trees with reversible DNA models, we used a partitioned model such that each partition was able to have an independent DNA substitution model. On the other hand, when we inferred the NR-DNA model we estimated a single model for the entire alignment. Thus, the NR-DNA model we inferred was unable to account for heterogeneity in the evolutionary process among partitions, possibly leading to its worse fit to the data when assessed using BIC scores. This suggests that using either mixture models or partitioned models may improve the fit of non-reversible DNA models to the data. The DNA results are consistent with results from previous study using the NR-DNA model and RootDigger (Bettisworth and Stamatakis 2020), although that study did not compare the performance of IQ-TREE and RootDigger on empirical datasets. Its results indicate that the NR-DNA model in IQ-TREE could not infer the correct root placement for any of the three tested datasets.

Our results demonstrate that the amino-acid non-reversible model can often be surprisingly accurate for inferring the root placement of phylogenies in the absence of additional information (such as outgroups) or assumptions (such as molecular clocks). In all of the well-defined clades that we examined, the non-reversible amino-acid model successfully identified the root that we identified a-priori as correct, and with very high rootstrap support. Importantly, the non-reversible amino-acid models also tended to fit the data far better than their reversible counterparts did. Indeed, we show that root placements appear to be accurate even with datasets as small as 50 well-curated loci between fairly closely-related taxa such as orders of mammals. Nevertheless, the application of the non-reversible amino acid models to two clades where the root position has previously been contentious failed to shed much additional light on the true root placement. Thus, while we show that the use of non-reversible models certainly has promise, we also show that it is no silver bullet.

Where a reliable outgroup taxon can be found, without the issues that can confound the inference of root placements using outgroups (Dalevi, et al. 2001; Braun and Kimball 2002; Graham, et al. 2002; Brady, et al. 2006), we suggest relying on the use of outgroups. Nevertheless, where no reliable outgroups can be found, or where there is some reason to question the position of a root inferred using an outgroup (e.g. references about questionable outgroup rooting), our study suggests that using non-reversible models can provide a useful additional line of evidence for the position of the root of a phylogeny. We note also that the rootstrap value and the AU test could be used to provide estimates of the uncertainty of root placement using an outgroup taxon Our work suggests a practical approach to inferring the root of a phylogenetic tree using non-reversible models. First, estimate an unrooted tree topology using the best reversible models available, excluding outgroup sequences. Next, fix the tree topology and use the best non-reversible models available to infer the Maximum Likelihood (ML) root position of that tree. Finally, determine to what extent the ML root position should be trusted. The degree of trust that researchers should put in an inferred ML root position should be influenced by three factors (noting of course that all phylogenetic inferences are susceptible to be misled by model misspecification). First, the fit of the non-reversible model to the data should be better than the fit of the reversible model. This can be assessed using common criteria like AICc or BIC scores. A better fit of the non-reversible model provides some assurance that the data contain sufficient signal that using a non-reversible model is advisable in the first place. Our results show that the root placement was inferred correctly with high rootstrap support in 12 out of the 13 datasets in which the non-reversible model was preferable.

In the absence of a better fit for a non-reversible model, we do not think any inferred ML root position should be trusted. Second, root positions with higher rootstrap support should be trusted more, because a higher rootstrap support indicates less variance among sites in the signal for the placement of the root. Third, ML root positions should be trusted more when the number of root placements included in the confidence set of an AU test is small, because a smaller confidence set indicates that there is less uncertainty in the root placement when the analysis is conditioned on the full alignment and the unrooted ML tree topology. A conservative approach to inferring root placements with non-reversible models would be to consider any root placement that has a substantial fraction of the rootstrap support and/or is included in the set of possible root placements identified by the AU test as a possible root placement given the assumptions of the model.

We hope that the combination of non-reversible, rootstrap support, and AU tests will add another tool to the phylogeneticist’s arsenal when it comes to inferring rooted phylogenies.

## References

Agnarsson I, Zambrana-Torrelio CM, Flores-Saldana NP, May-Collado LJ. 2011. A time-calibrated species-level phylogeny of bats (Chiroptera, Mammalia). PLoS Curr 3:RRN1212.

Allman ES, Degnan JH, Rhodes JA. 2011. Identifying the rooted species tree from the distribution of unrooted gene trees under the coalescent. J. Math. Biol. 62:833–862.

Baldauf SL, Palmer JD. 1993. Animals and fungi are each other’s closest relatives: congruent evidence from multiple proteins. Proceedings of the National Academy of Sciences 90:11558-11562.

Bettisworth B, Stamatakis A. 2020. RootDigger: a root placement program for phylogenetic trees. bioRxiv:2020.2002.2013.935304.

Boussau B, Szollosi GJ, Duret L, Gouy M, Tannier E, Daubin V. 2013. Genome-scale coestimation of species and gene trees. Genome Res. 23:323-330.

Brady SG, Schultz TR, Fisher BL, Ward PS. 2006. Evaluating alternative hypotheses for the early evolution and diversification of ants. Proc Natl Acad Sci U S A 103:18172–18177.

Braun EL, Kimball RT. 2002. Examining Basal avian divergences with mitochondrial sequences: model complexity, taxon sampling, and sequence length. Syst. Biol. 51:614–625.

Cherlin S, Heaps SE, Nye TMW, Boys RJ, Williams TA, Embley TM. 2018. The Effect of Nonreversibility on Inferring Rooted Phylogenies. Mol. Biol. Evol. 35:984–1002.

Chernomor O, von Haeseler A, Minh BQ. 2016. Terrace Aware Data Structure for Phylogenomic Inference from Supermatrices. Syst. Biol. 65:997–1008.

Dalevi D, Hugenholtz P, Blackall LL. 2001. A multiple-outgroup approach to resolving division-level phylogenetic relationships using 16S rDNA data. Int. J. Syst. Evol. Microbiol. 51:385–391.

Dayhoff M. 1987. A model of evolutionary change in proteins. Atlas of protein sequence and structure 5:suppl. 3.

Dayhoff M, Schwartz R. 1980. Prokaryote evolution and the symbiotic origin of eukaryotes. Endocytobiology: endosymbiosis and cell biology: a synthesis of recent research 1:63–84.

Dayhoff M, Schwartz R, Orcutt B. 1978. A model of evolutionary change in proteins. Atlas of protein sequence and structure 5:345-352.

Drummond AJ, Ho SY, Phillips MJ, Rambaut A. 2006. Relaxed phylogenetics and dating with confidence. PLoS Biol. 4:e88.

Duchene DA, Tong KJ, Foster CSP, Duchene S, Lanfear R, Ho SYW. 2020. Linking Branch Lengths across Sets of Loci Provides the Highest Statistical Support for Phylogenetic Inference. Mol. Biol. Evol. 37:1202–1210.

Farris JS. 1972. Estimating Phylogenetic Trees from Distance Matrices. Am. Nat. 106:645-&.

Gatesy J, Springer MS. 2017. Phylogenomic red flags: Homology errors and zombie lineages in the evolutionary diversification of placental mammals. Proc Natl Acad Sci U S A 114:E9431-E9432.

Gogarten JP, Kibak H, Dittrich P, Taiz L, Bowman EJ, Bowman BJ, Manolson MF, Poole RJ, Date T, Oshima T, et al. 1989. Evolution of the vacuolar H+-ATPase: implications for the origin of eukaryotes. Proc Natl Acad Sci U S A 86:6661-6665.

Graham SW, Olmstead RG, Barrett SC. 2002. Rooting phylogenetic trees with distant outgroups: a case study from the commelinoid monocots. Mol. Biol. Evol. 19:1769-1781.

Hasegawa M, Kishino H, Yano T. 1985. Dating of the human-ape splitting by a molecular clock of mitochondrial DNA. J. Mol. Evol. 22:160-174.

Heaps SE, Nye TM, Boys RJ, Williams TA, Embley TM. 2014. Bayesian modelling of compositional heterogeneity in molecular phylogenetics. Stat Appl Genet Mol Biol 13:589-609.

Holmes S. 2003. Bootstrapping phylogenetic trees: Theory and methods. Statistical Science 18:241-255.

Huelsenbeck JP, Bollback JP, Levine AM. 2002. Inferring the Root of a Phylogenetic Tree. Syst. Biol. 51:32-43.

Iwabe N, Kuma K, Hasegawa M, Osawa S, Miyata T. 1989. Evolutionary relationship of archaebacteria, eubacteria, and eukaryotes inferred from phylogenetic trees of duplicated genes. Proc Natl Acad Sci U S A 86:9355-9359.

Jansen RK, Cai Z, Raubeson LA, Daniell H, Depamphilis CW, Leebens-Mack J, Muller KF, Guisinger-Bellian M, Haberle RC, Hansen AK, et al. 2007. Analysis of 81 genes from 64 plastid genomes resolves relationships in angiosperms and identifies genome-scale evolutionary patterns. Proc Natl Acad Sci U S A 104:19369-19374.

Jayaswal V, Ababneh F, Jermiin LS, Robinson J. 2011. Reducing model complexity of the general Markov model of evolution. Mol. Biol. Evol. 28:3045-3059.

Jones DT, Taylor WR, Thornton JM. 1992. The rapid generation of mutation data matrices from protein sequences. Comput. Appl. Biosci. 8:275-282.

Jukes TH, Cantor C. 1969. Evolution of protein molecules. In: Munro HN, editor. In Mammalian Protein Metabolism.

Kalyaanamoorthy S, Minh BQ, Wong TKF, von Haeseler A, Jermiin LS. 2017. ModelFinder: fast model selection for accurate phylogenetic estimates. Nat. Methods 14:587-589.

Kimura M. 1980. A Simple Method for Estimating Evolutionary Rates of Base Substitutions through Comparative Studies of Nucleotide-Sequences. J. Mol. Evol. 16:111-120.

Kocot KM, Cannon JT, Todt C, Citarella MR, Kohn AB, Meyer A, Santos SR, Schander C, Moroz LL, Lieb B, et al. 2011. Phylogenomics reveals deep molluscan relationships. Nature 477:452-456.

Lake JA, Herbold CW, Rivera MC, Servin JA, Skophammer RG. 2007. Rooting the tree of life using nonubiquitous genes. Mol. Biol. Evol. 24:130-136.

Le SQ, Gascuel O. 2008. An improved general amino acid replacement matrix. Mol. Biol. Evol. 25:1307-1320.

Liu L, Zhang J, Rheindt FE, Lei F, Qu Y, Wang Y, Zhang Y, Sullivan C, Nie W, Wang J, et al. 2017. Genomic evidence reveals a radiation of placental mammals uninterrupted by the KPg boundary. Proc Natl Acad Sci U S A 114:E7282-E7290.

Lyons-Weiler J, Hoelzer GA, Tausch RJ. 1998. Optimal outgroup analysis. Biol. J. Linn. Soc. 64:493-511.

Maddison WP, Donoghue MJ, Maddison DR. 1984. Outgroup Analysis and Parsimony. Systematic Zoology 33:83-103.

Mai U, Sayyari E, Mirarab S. 2017. Minimum variance rooting of phylogenetic trees and implications for species tree reconstruction. PLoS One 12:e0182238.

Meganathan PR, Pagan HJ, McCulloch ES, Stevens RD, Ray DA. 2012. Complete mitochondrial genome sequences of three bats species and whole genome mitochondrial analyses reveal patterns of codon bias and lend support to a basal split in Chiroptera. Gene 492:121-129.

Meredith RW, Janecka JE, Gatesy J, Ryder OA, Fisher CA, Teeling EC, Goodbla A, Eizirik E, Simao TL, Stadler T, et al. 2011. Impacts of the Cretaceous Terrestrial Revolution and KPg extinction on mammal diversification. Science 334:521-524.

Milinkovitch MC, Lyons-Weiler J. 1998. Finding optimal ingroup topologies and convexities when the choice of outgroups is not obvious. Mol. Phylogen. Evol. 9:348-357.

Minh BQ, Hahn MW, Lanfear R. 2020. New methods to calculate concordance factors for phylogenomic datasets. Mol. Biol. Evol.

Minh BQ, Schmidt HA, Chernomor O, Schrempf D, Woodhams MD, von Haeseler A, Lanfear R. 2020. IQ-TREE 2: New Models and Efficient Methods for Phylogenetic Inference in the Genomic Era. Mol. Biol. Evol. 37:1530-1534.

Moore MJ, Bell CD, Soltis PS, Soltis DE. 2007. Using plastid genome-scale data to resolve enigmatic relationships among basal angiosperms. Proc Natl Acad Sci U S A 104:19363-19368.

Nardi F, Spinsanti G, Boore JL, Carapelli A, Dallai R, Frati F. 2003. Hexapod origins: monophyletic or paraphyletic? Science 299:1887-1889.

Naser-Khdour S, Minh BQ, Zhang W, Stone EA, Lanfear R. 2019. The Prevalence and Impact of Model Violations in Phylogenetic Analysis. Genome Biol Evol.

Philippe H, Brinkmann H, Lavrov DV, Littlewood DT, Manuel M, Worheide G, Baurain D. 2011. Resolving difficult phylogenetic questions: why more sequences are not enough. PLoS Biol. 9:e1000602.

Ren M, Sun HJ, Bo SQ, Zhang SY, Hua PY. 2018. Parallel amino acid deletions of prestin protein in two dramatically divergent bat lineages suggest the complexity of the evolution of echolocation in bats. Acta Chiropterologica 20:311-317.

Reyes-Amaya N, Flores D. 2019. Hypophysis size evolution in Chiroptera. Acta Chiropterologica 21:65-74.

Rivera MC, Lake JA. 1992. Evidence That Eukaryotes and Eocyte Prokaryotes Are Immediate Relatives. Science 257:74-76.

Shen XX, Hittinger CT, Rokas A. 2017. Contentious relationships in phylogenomic studies can be driven by a handful of genes. Nat Ecol Evol 1:126.

Shimodaira H. 2002. An approximately unbiased test of phylogenetic tree selection. Syst. Biol. 51:492-508.

Shimodaira H, Hasegawa M. 1999. Multiple comparisons of log-likelihoods with applications to phylogenetic inference. Mol. Biol. Evol. 16:1114-1116.

Smith AB. 1994. Rooting Molecular Trees - Problems and Strategies. Biol. J. Linn. Soc. 51:279-292.

Squartini F, Arndt PF. 2008. Quantifying the Stationarity and Time Reversibility of the Nucleotide Substitution Process. Molecular Biology and Evolution 25:2525-2535.

Steenkamp ET, Wright J, Baldauf SL. 2006. The protistan origins of animals and fungi. Mol. Biol. Evol. 23:93-106.

Swofford D, Olsen G, Waddell P. 1996. Phylogenetic inference. In: David M. Hillis CM, Barbara K. Mable, editor. Molecular Systematics, 2nd edn: Sunderland, Mass. : Sinauer Associates. p. 407-513.

Tamura K, Nei M. 1993. Estimation of the number of nucleotide substitutions in the control region of mitochondrial DNA in humans and chimpanzees. Mol. Biol. Evol. 10:512-526.

Tavaré S. 1986. Some probabilistic and statistical probles in the analysis of DNA sequences. Lectures on Mathematics in the Life Sciences 17.

Tria FDK, Landan G, Dagan T. 2017. Phylogenetic rooting using minimal ancestor deviation. Nat Ecol Evol 1:193.

Tsagkogeorga G, Parker J, Stupka E, Cotton JA, Rossiter SJ. 2013. Phylogenomic analyses elucidate the evolutionary relationships of bats. Curr. Biol. 23:2262-2267.

Watrous LE, Wheeler QD. 1981. The out-Group Comparison Method of Character Analysis. Systematic Zoology 30:1-11.

Whelan NV, Kocot KM, Moroz LL, Halanych KM. 2015. Error, signal, and the placement of Ctenophora sister to all other animals. Proc Natl Acad Sci U S A 112:5773-5778.

Whelan S, Goldman N. 2001. A general empirical model of protein evolution derived from multiple protein families using a maximum-likelihood approach. Mol. Biol. Evol. 18:691-699.

Williams KP, Gillespie JJ, Sobral BW, Nordberg EK, Snyder EE, Shallom JM, Dickerman AW. 2010. Phylogeny of gammaproteobacteria. J. Bacteriol. 192:2305-2314.

Williams TA, Heaps SE, Cherlin S, Nye TM, Boys RJ, Embley TM. 2015. New substitution models for rooting phylogenetic trees. Philos Trans R Soc Lond B Biol Sci 370:20140336.

Wu S, Edwards S, Liu L. 2019. Data from: Genome-scale DNA sequence data and the evolutionary history of placental mammals. In: Figshare.

Wu S, Edwards S, Liu L. 2018. Genome-scale DNA sequence data and the evolutionary history of placental mammals. Data Brief 18:1972-1975.

Yang ZH, Roberts D. 1995. On the Use of Nucleic-Acid Sequences to Infer Early Branchings in the Tree of Life. Mol. Biol. Evol. 12:451-458.

Yap VB, Speed T. 2005. Rooting a phylogenetic tree with nonreversible substitution models. BMC Evol. Biol. 5:2.

Yu Y, Warnow T, Nakhleh L. 2011. Algorithms for MDC-based multi-locus phylogeny inference: beyond rooted binary gene trees on single alleles. J. Comput. Biol. 18:1543-1559.

Zhang SQ, Che LH, Li Y, Dan L, Pang H, Slipinski A, Zhang P. 2018. Evolutionary history of Coleoptera revealed by extensive sampling of genes and species. Nat Commun 9:205.

Zhou X, Xu S, Yang Y, Zhou K, Yang G. 2011. Phylogenomic analyses and improved resolution of Cetartiodactyla. Mol Phylogenet Evol 61:255-264.

