## Appendix for "Assessing Confidence in Root Placement on Phylogenies: An Empirical Study Using Non-Reversible Models for Mammals"

### ALGORITHM A.1.

Since we want to define the well-defined clades for the non-reversible analysis based on the bootstrap support values and the concordance factors values we will use the following algorithm to find well-defined clades in the whole MSA:

(1) First, estimate the unrooted topology and the bootstrap support values using the ultrafast bootstrap (UFBoot) with 1000 replicates (Hoang, et al. 2018)

```
iqtree2 –s ALIGNMENT_FILE –p PARTITION_FILE –B 1000 --prefix REV
```

Where ALIGNMENT_FILE is the MSA file, PARTITION_FILE is the partition file (it can be the same as the MSA file), -p is the option for edge-linked substitution rates,–B is option for UFBoot with 1000 replicates, and --prefix so all the output files will be named REV.*.

(2) Infer the single-locus trees

```
iqtree2 –s ALIGNMENT_FILE –S PARTITION_FILE --prefix LOCI
```

Like –p option, the –S option performs model selection for each loci separately. However, unlike –p option, the –S option infer a separate tree for each loci. All output files are of the form LOCI.*.

(3) calculate the gCF and sCF (Minh, et al. 2018) for every branch of the ML species tree

```
Iqtree2 –t REV.treefile --gcf LOCI.treefile –s ALIGNMENT_FILE --scf 100 --prefix CONCORD
```

Where REV.treefile is the unrooted species tree, and LOCI.treefile is file with all the unrooted loci trees.

**ALGORITHM A.2.**

For each sub-dataset that we want to infer the root placement using the non-reversible model. However, as the non-reversible model is computationally expensive, we first find the best partitioning scheme using the reversible model and then use this scheme to find the non-reversible model parameters and infer the rooted tree.

(1) Find the best partitioning scheme using the best-fit reversible model

```
iqtree2 -s ALIGNMENT_FILE -p PARTITION_FILE --prefix REV
```

(2) Estimate the non-reversible models' parameters using the best partitioning scheme from the previous step and unrooted tree as the initial tree

For NR-AA:

```
iqtree2 -s ALIGNMENT_FILE -p REV.best_scheme.nex -t REV.treefile
--model-joint NONREV -B 1000 --prefix AA_NONREV
```

For NR-DNA:

```
iqtree2 -s ALIGNMENT_FILE -p REV.best_scheme.nex -t REV.treefile
--model-joint 12.12 -B 1000 --prefix DNA_NONREV
```

Where NONREV and 12.12 are the substitution models for NR-AA and NR-DNA as defined in IQ-TREE, respectively.

(3) Compare the BIC scores of the reversible and non-reversible models.

If $BIC_{NONREV} > BIC_{REV}$, then using non-reversible models to infer the rooted topology is not advised due to over-parameterization of the data. In this case, an alternative rooting method is recommended if possible.

### ALGORITHM A.3.

We apply the AU test (Shimodaira 2002) to all the possible rooting placements using the following command-line:

For NR-AA:

```
iqtree2 –s ALIGNMENT_FILE –p REV.best_scheme.nex –model-joint NONREV
--root-test –zb 1000 –au –te NONREV.treefile --prefix TOP
```

For NR-DNA:

```
iqtree2 –s ALIGNMENT_FILE –p REV.best_scheme.nex –model-joint 12.12
--root-test –zb 1000 –au –te NONREV.treefile --prefix TOP
```

Where --root-test will re-root the tree on every branch, -zb option for specifying the number of RELL replicates (Kishino, et al. 1990), and -au option to perform the AU test.

### TABLES

Table A.1. Last decade's relevant literature of the well-defined clades that we used in this study for estimating the root.

| Clade | Study Reference | Root placement |
|---|---|---|
| Afrotheria | (Elliot and Crespi 2009) | {Afroinsectiphilia, Paenungulata} |
| | (Asher, et al. 2009) | {Afroinsectiphilia, Paenungulata} |
| | (Poulakakis and Stamatakis 2010) | {Afroinsectiphilia, Paenungulata} |
| | (Phillips and Penny 2010) | {Afroinsectiphilia, Paenungulata} |
| | (Romiguier, et al. 2010) | {Afroinsectiphilia, Paenungulata} |
| | (Kuntner, et al. 2011) | {Afroinsectiphilia, Paenungulata} |
| | (Meredith, et al. 2011) | {Afroinsectiphilia, Paenungulata} |
| | (Svartman and Stanyon 2012) | {Afroinsectiphilia, Paenungulata} |
| | (Lartillot and Delsuc 2012) | {Afroinsectiphilia, Paenungulata} |
| | (dos Reis, et al. 2012) | {Afroinsectiphilia, Paenungulata} |
| | (Benoit, et al. 2013) | {Afroinsectiphilia, Paenungulata} |
| | (Wu, et al. 2014) | {Afroinsectiphilia, Paenungulata} |
| | (Gheerbrant, et al. 2014) | {Afroinsectiphilia, Paenungulata} |
| | (Halliday, et al. 2015) | {Afroinsectiphilia, Paenungulata} |
| | (Puttick and Thomas 2015) | {Afrosoricida },{Afroinsectiphilia, Paenungulata} |
| | (Foley, et al. 2016) | {Afroinsectiphilia, Paenungulata} |
| | (Liu, et al. 2017) | {Afroinsectiphilia, Paenungulata} |
| | (Wu, et al. 2017) | {Afroinsectiphilia, Paenungulata} |
| Primates | (Fabre, et al. 2009) | {Strepsirrhini, Haplorrhini} |
| | (Chatterjee, et al. 2009) | {Strepsirrhini, Haplorrhini} |
| | (Matsui, et al. 2009) | {Trasiiformes},{Strepsirrhini, Haplorrhini} |
| | (Romiguier, et al. 2010) | {Strepsirrhini, Haplorrhini} |
| | (Perelman, et al. 2011) | {Strepsirrhini, Haplorrhini} |
| | (Meredith, et al. 2011) | {Strepsirrhini, Haplorrhini} |
| | (Jameson, et al. 2011) | {Strepsirrhini, Haplorrhini} |
| | (Diogo and Wood 2011) | {Strepsirrhini, Haplorrhini} |
| | (Springer, et al. 2012) | {Strepsirrhini, Haplorrhini} |
| | (dos Reis, et al. 2012) | {Strepsirrhini, Haplorrhini} |
| | (Steiper and Seiffert 2012) | {Strepsirrhini, Haplorrhini} |
| | (Lartillot and Delsuc 2012) | {Strepsirrhini, Haplorrhini} |
| | (Finstermeier, et al. 2013) | {Strepsirrhini, Haplorrhini} |
| | (Hartig, et al. 2013) | {Strepsirrhini, Haplorrhini} |
| | (Kumar, et al. 2013) | {Strepsirrhini, Haplorrhini} |
| | (Pozzi, et al. 2014) | {Strepsirrhini, Haplorrhini} |
| | (Wu, et al. 2014) | {Strepsirrhini, Haplorrhini} |
| | (Pattinson, et al. 2015) | {Strepsirrhini, Haplorrhini} |

| | | |
|---|---|---|
| | (Kari, et al. 2015) | {Strepsirrhini, Haplorrhini} |
| | (Herrera and Davalos 2016) | {Strepsirrhini, Haplorrhini} |
| | (Liu, et al. 2017) | {Strepsirrhini, Haplorrhini} |
| | (Wu, et al. 2017) | {Strepsirrhini, Haplorrhini} |
| | (Monson and Hlusko 2018) | {Strepsirrhini, Haplorrhini} |
| | (Reis, et al. 2018) | {Strepsirrhini, Haplorrhini} |
| | (Zhang, et al. 2019) | {Strepsirrhini, Haplorrhini} |
| Myomorpha | (Blanga-Kanfi, et al. 2009) | {Muroidea, Dipodoidea} |
| | (Churakov, et al. 2010) | {Muroidea, Dipodoidea} |
| | (Hao, et al. 2011) | {Muroidea, Dipodoidea} |
| | (Meredith, et al. 2011) | {Muroidea, Dipodoidea} |
| | (Horn, et al. 2011) | {Muroidea, Dipodoidea} |
| | (Fabre, et al. 2012) | {Muroidea, Dipodoidea} |
| | (Wu, et al. 2012) | {Muroidea, Dipodoidea} |
| | (Schenk, et al. 2013) | {Muroidea, Dipodoidea} |
| | (Wu, et al. 2014) | {Muroidea, Dipodoidea} |
| | (Yue, et al. 2015) | {Muroidea, Dipodoidea} |
| | (Liu, et al. 2017) | {Muroidea, Dipodoidea} |
| | (Wu, et al. 2017) | {Muroidea, Dipodoidea} |
| | (Tavares and Seuanez 2018) | {Muroidea, Dipodoidea} |
| | (Swanson, et al. 2019) | {Muroidea, Dipodoidea} |
| | (Hedrick, et al. 2020) | {Muroidea, Dipodoidea} |
| Carnivora | (Finarelli and Flynn 2009) | {Feliformia,Caniformia} |
| | (Agnarsson, et al. 2010) | {Feliformia,Caniformia} |
| | (Eizirik, et al. 2010) | {Feliformia,Caniformia} |
| | (Stankowich, et al. 2011) | {Feliformia,Caniformia} |
| | (Nyakatura and Bininda-Emonds 2012) | {Feliformia,Caniformia} |
| | (Lartillot and Delsuc 2012) | {Feliformia,Caniformia} |
| | (dos Reis, et al. 2012) | {Feliformia,Caniformia} |
| | (Wu, et al. 2014) | {Feliformia,Caniformia} |
| | (Tomiya and Tseng 2016) | {Feliformia,Caniformia} |
| | (Panciroli, et al. 2017) | {Feliformia,Caniformia} |
| | (Liu, et al. 2017) | {Feliformia,Caniformia} |
| | (Wu, et al. 2017) | {Feliformia,Caniformia} |
| | (Polly, et al. 2017) | {Feliformia,Caniformia} |
| | (Machado, et al. 2018) | {Feliformia,Caniformia} |
| Bovidae | (Bibi, et al. 2009) | {Bovinae} |
| | (Hassanin, et al. 2012) | {Bovinae} |
| | (Yang, et al. 2013) | {Bovinae} |
| | (Wu, et al. 2014) | {Bovinae} |
| | (Bibi 2013) | {Bovinae} |
| | (Liu, et al. 2017) | {Bovinae} |

| | |
|---|---|
| (Wu, et al. 2017) | {Bovinae} |
| (Chen, et al. 2019) | {Bovinae} |

Table A.2. The well-defined clades that we used in this study for estimating the root.

| Clade | No. taxa | Dataset | branch | %BS | gCF | gDF1 | gDF2 | sCF |
|---|---|---|---|---|---|---|---|---|
| Afrotheria | 7 | AA | leading | 100.0 | 55.18 | 0.32 | 0.66 | 55.34 |
| | | | descen.1 | 100.0 | 24.68 | 5.38 | 8.30 | 35.64 |
| | | | descen.2 | 100.0 | 55.43 | 1.41 | 1.01 | 55.12 |
| | | DNA | leading | 100.0 | 52.72 | 0.18 | 0.42 | 50.56 |
| | | | descen.1 | 100.0 | 17.97 | 6.23 | 8.07 | 32.87 |
| | | | descen.2 | 100.0 | 50.95 | 0.96 | 1.41 | 51.22 |
| Primates | 16 | AA | leading | 100.0 | 25.10 | 2.14 | 2.67 | 41.22 |
| | | | descen.1 | 100.0 | 25.77 | 8.82 | 7.91 | 37.91 |
| | | | descen.2 | 100.0 | 45.10 | 1.25 | 1.75 | 55.00 |
| | | DNA | leading | 100.0 | 28.04 | 2.19 | 2.79 | 38.79 |
| | | | descen.1 | 100.0 | 27.29 | 9.29 | 7.63 | 35.82 |
| | | | descen.2 | 100.0 | 47.64 | 1.42 | 1.67 | 50.39 |
| Myomorpha | 7 | AA | leading | 100.0 | 58.10 | 5.94 | 5.9 | 48.09 |
| | | | descen.1 | 100.0 | 84.47 | 0.79 | 0.61 | 73.88 |
| | | DNA | leading | 100.0 | 57.47 | 5.36 | 5.74 | 43.32 |
| | | | descen.1 | 100.0 | 83.95 | 0.62 | 0.50 | 65.66 |
| Carnivora | 9 | AA | leading | 100.0 | 60.22 | 1.01 | 1.51 | 64.41 |
| | | | descen.1 | 100.0 | 82.06 | 0.53 | 0.76 | 94.15 |
| | | | descen.2 | 100.0 | 49.80 | 5.62 | 7.87 | 50.53 |
| | | DNA | leading | 100.0 | 53.63 | 6.11 | 6.51 | 45.8 |
| | | | descen.1 | 100.0 | 84.23 | 0.50 | 0.62 | 92.34 |
| | | | descen.2 | 100.0 | 51.18 | 3.88 | 6.2 | 50.73 |
| Bovidae | 5 | AA | leading | 100.0 | 85.66 | 0.18 | 0.16 | 89.47 |
| | | | descen.1 | 100.0 | 81.28 | 3.9 | 2.38 | 93.25 |
| | | | descen.2 | 100.0 | 71.83 | 5.71 | 5.28 | 79.17 |
| | | DNA | leading | 100.0 | 85.53 | 0.28 | 0.21 | 85.37 |
| | | | descen.1 | 100.0 | 81.78 | 3.64 | 2.27 | 90.19 |
| | | | descen.2 | 100.0 | 72.21 | 5.38 | 4.97 | 70.74 |

Note: The bootstrap, gCF, gDF1, gDF2, and sCF values are for the branch leading to that clade and the first direct descendants of the clade, respectively.

TABLE A.3. Rootstrap support value, number of alternative root placements in the AU test confidence set, rBED, and rSED for the amino-acid rooted trees of each clade.

| Clade | #loci | #sites | %RS support for true root | Is true root in CS? | root placements in CS | rBED | rSED |
|---|---|---|---|---|---|---|---|
| Carnivora | 5,162 | 3,050,199 | 100% | Yes | 0 | 0.0000 – 0.0273 | 0 |
| | 516 | 295,962 | 99.8% | Yes | 0 | 0.0000 – 0.0274 | 0 |
| | 52 | 33,404 | 99.1% | Yes | 0 | 0.0000 – 0.0242 | 0 |
| | 5 | 2,789 | 4.6% | No | 1 | 0.0074 – 0.0740 | 1 |
| Bovidae | 5,162 | 3,050,199 | 100% | Yes | 0 | 0.0000 – 0.0128 | 0 |
| | 516 | 310,353 | 100% | Yes | 0 | 0.0000 – 0.0130 | 0 |
| | 52 | 28,729 | 99.9% | Yes | 0 | 0.0000 – 0.0092 | 0 |
| | 5 | 3,222 | 41.9% | Yes | 0 | 0.0000 – 0.0210 | 0 |
| Myomorpha | 5,162 | 3,050,199 | 100.0% | Yes | 0 | 0.0000 – 0.0842 | 0 |
| | 516 | 309,459 | 100.0% | Yes | 0 | 0.0000 – 0.0886 | 0 |
| | 52 | 35,494 | 100.0% | Yes | 0 | 0.0000 – 0.0679 | 0 |
| | 5 | 4,438 | 36.6% | Yes | 0 | 0.0000 – 0.1135 | 0 |
| Primates | 5,162 | 3,050,199 | 100.0% | Yes | 0 | 0.0000 – 0.0092 | 0 |
| | 516 | 302,781 | 87.9% | Yes | 0 | 0.0000 – 0.0090 | 0 |
| | 52 | 30,225 | 59.8% | Yes | 0 | 0.0000 – 0.0106 | 0 |
| | 5 | 4,426 | 66.1% | Yes | 0 | 0.0000 – 0.0076 | 0 |
| Afrotheria | 5,162 | 3,050,199 | 100.0% | Yes | 0 | 0.0000 – 0.0077 | 0 |
| | 516 | 292,632 | 98.5% | Yes | 0 | 0.0000 – 0.0066 | 0 |
| | 52 | 28,904 | 29.2% | No | 1 | 0.0004 – 0.0127 | 1 |
| | 5 | 3,265 | 62.1% | Yes | 0 | 0.0000 – 0.0120 | 0 |

TABLE A.4. Rootstrap support value, number of alternative root placements in the AU test confidence set, rBED, and rSED for the nucleotide rooted trees of each clade.

| Clade | #loci | #sites | %RS support for true root | Is true root in CS? | root placements in CS | rBED | rSED |
|---|---|---|---|---|---|---|---|
| Carnivora | 15,486 | 9,150,597 | 100% | Yes | 0 | 0.0000 – 0.0229 | 0 |
| | 1,548 | 917,265 | 100% | Yes | 1 | 0.0000 – 0.0273 | 0 |
| | 154 | 99,334 | 99.1% | Yes | 0 | 0.0000 – 0.0238 | 0 |
| | 15 | 11,197 | 17.5% | No | 1 | 0.0199 – 0.0420 | 1 |
| Bovidae | 15,486 | 9,150,597 | 100.0% | Yes | 0 | 0.0000 – 0.0269 | 0 |
| | 1,548 | 910,022 | 100.0% | Yes | 0 | 0.0000 – 0.0269 | 0 |
| | 154 | 91,674 | 40.8% | Yes | 0 | 0.0000 – 0.0196 | 0 |
| | 15 | 5,560 | 1.4% | No | 2 | 0.0036 – 0.0198 | 2 |
| Myomorpha | 15,485 | 9,149,793 | 73.2% | Yes | 1 | 0.0000 – 0.1292 | 0 |
| | 1,548 | 898,643 | 25.9% | Yes | 1 | 0.0048 – 0.0150 | 1 |
| | 154 | 87,033 | 5.4% | Yes | 1 | 0.0159 – 0.0253 | 1 |
| | 15 | 9,433 | 15.7% | No | 1 | 0.0352 – 0.0431 | 2 |
| Primates | 15,486 | 9,150,597 | 99.7% | Yes | 0 | 0.0000 – 0.0096 | 0 |
| | 1,548 | 912,110 | 87.0% | Yes | 0 | 0.0000 – 0.0087 | 0 |
| | 154 | 93,830 | 31.1% | Yes | 0 | 0.0132 – 0.0300 | 1 |
| | 15 | 8,242 | 3.0% | Yes | 1 | 0.0176 – 0.0221 | 4 |
| Afrotheria | 15,486 | 9,150,597 | 0.0% | No | 1 | 0.0157 – 0.0268 | 2 |
| | 1,548 | 924,981 | 21.0% | No | 1 | 0.0133 – 0.0251 | 2 |
| | 154 | 92,545 | 44.4% | No | 1 | 0.0000 – 0.0073 | 0 |
| | 15 | 6,697 | 4.7% | No | 1 | 0.0268 – 0.0428 | 2 |

TABLE A.5. Number of sites, root placement, RS support for the ML root placement, better BIC score (between R and NR models), and root placements in the AU confidence set for Chiroptera.

| | dataset | #sites | Root placement | RS% ML root | Better BIC | AU CS |
|---|---|---|---|---|---|---|
| DNA-NR | Whole dataset | 9,149,793 | Y-Y | 100.0% | R | {Y-Y} |
| | Subsampled 10% | 921,910 | Y-Y | 99.9% | R | {Y-Y} |
| | Subsampled 1% | 85,437 | Y-Y | 41.6% | R | {Y-Y} |
| | Subsampled 0.1% | 7,804 | Y-Y | 9.6% | R | {Y-Y} |
| | MaxSym test | 6,854,459 | Y-Y | 90.1% | NR | {Y-Y} |
| | Highest GLS | 8,769,211 | Y-Y | 100.0% | R | {Y-Y} |
| AA-NR | Whole dataset | 3,050,199 | Y-Y | 65.5% | NR | {Y-Y, M-M, R, P} |
| | Subsampled 10% | 309,745 | Y-Y | 51.0% | NR | {Y-Y, M-M} |
| | Subsampled 1% | 33,365 | R | 38.1% | R | {Y-Y, R, P} |
| | Subsampled 0.1% | 3,955 | M-M | 70.2% | R | {Y-Y, M-M} |
| | MaxSym test | 2,934,731 | Y-Y | 65.9% | NR | {Y-Y, M-M, R } |
| | Highest GLS | 3,027,379 | Y-Y | 76.6% | NR | {Y-Y, M-M} |

Note: Y-Y refers to Yinptero-Yango hypothesis, M-M refers to Micro-Mega hypothesis, R refers to Rhinolophoidea and P refers to Pteropus (Flying foxes). Highest GLS refers to dataset where the top 1% loci with the highest ΔGLS removed.

TABLE A.6. Number of sites, root placement, RS support for the ML root placement, better BIC score (between R and NR models, and root placements in the AU confidence set for Cetartiodactyla.

| | dataset | #sites | Root placement | RS% ML root | Better AICc | AU CS |
|---|---|---|---|---|---|---|
| DNA-NR | Whole dataset | 9,149,793 | T | 71.0% | R | {T} |
| | Subsampled 10% | 915,274 | T | 84.5% | R | {T} |
| | Subsampled 1% | 90,371 | T+S | 38.9% | R | {S} |
| | Subsampled 0.1% | 6,003 | T | 41.6% | R | {T} |
| | MaxSym test | 7,565,809 | T | 68.7% | R | {T} |
| | Highest GLS | 8,557,372 | T+S | 39.7% | R | {S} |
| AA-NR | Whole dataset | 3,050,199 | T+S | 71.8% | NR | {T+S, T, S} |
| | Subsampled 10% | 293,123 | T | 62.9% | NR | {T+S} |
| | Subsampled 1% | 31,253 | T+S | 50.2% | R | {T+S, T} |
| | Subsampled 0.1% | 1,480 | B | 40.7% | R | {T+S, T} |

| | | | | | |
|---|---|---|---|---|---|
| MaxSym test | 2,997,304 | T+S | 63.6% | NR | {T+S, T} |
| Highest GLS | 3,024,886 | T+S | 63.2% | NR | {T+S, T} |

Note: T refers to Tylopoda, S refers to Suina, and B refers to Bovidae. Highest GLS refers to dataset where the top 1% loci with the highest ΔGLS removed.

TABLE A.7. Likelihood, number of free parameters, and BIC score of the amino-acid reversible and non-reversible models.

| Clade | #loci | Rev-AA | | | NR-AA | | |
|---|---|---|---|---|---|---|---|
| | | logL | #FP | BIC | logL | #FP | BIC |
| Afrotheria | 5,162 | -14330208.59 | 21172 | 28976530 | -14297752.78 | 9716 | 28740572 |
| | 516 | -1357177.681 | 2117 | 2741001 | -1354652.159 | 1320 | 2725919 |
| | 52 | -136982.9062 | 220 | 276226 | -136452.7464 | 487 | 277908 |
| | 5 | -19001.63 | 58 | 38473 | -18665.2888 | 401 | 40575 |
| Bovidae | 5,162 | -9495698.792 | 16456 | 19237097 | -9526590.714 | 6026 | 19143154 |
| | 516 | -969922.7367 | 1716 | 1961545 | -972933.315 | 957 | 1957968 |
| | 52 | -90113.4964 | 100 | 181254 | -90307.5227 | 443 | 185163 |
| | 5 | -10558.2498 | 11 | 21205 | -10429.3843 | 392 | 24025 |
| Carnivora | 5,162 | -11771808.39 | 19563 | 23835706 | -11783521.78 | 8525 | 23694328 |
| | 516 | -1159792.644 | 1907 | 2343610 | -1160442.419 | 1205 | 2336065 |
| | 52 | -133619.2048 | 288 | 270238 | -133736.2131 | 479 | 272462 |
| | 5 | -12401.6075 | 37 | 25097 | -12212.3584 | 399 | 27590 |
| Myomorpha | 5,162 | -13717877.69 | 21337 | 27754332 | -13704191.03 | 9710 | 27553359 |
| | 516 | -1410118.562 | 2041 | 2846041 | -1407094.056 | 1320 | 2830876 |
| | 52 | -159843.2064 | 219 | 321981 | -159218.9652 | 486 | 323530 |
| | 5 | -19717.5842 | 38 | 39754 | -19553.1957 | 400 | 42466 |
| Primates | 5,162 | -14060075.69 | 19597 | 28412749 | -14035582.5 | 9433 | 28212006 |
| | 516 | -1409968.454 | 1965 | 2844737 | -1406902.357 | 1320 | 2830464 |

| | | | | | | |
|---|---|---|---|---|---|---|
| 52 | -135178.7103 | 209 | 272514 | -134377.0976 | 495 | 273861 |
| 5 | -18618.1041 | 74 | 37857 | -18440.361 | 417 | 40382 |

TABLE A.8. Likelihood, number of free parameters, and BIC score of nucleotide reversible and non-reversible models.

| | | Rev-DNA | | | NR-DNA | | |
|---|---|---|---|---|---|---|---|
| Clade | #loci | logL | #FP | BIC | logL | #FP | BIC |
| Afrotheria | 15,486 | -25649218.58 | 83714 | 52640316 | -26103942.35 | 26592 | 52634137 |
| | 1,548 | -2592184.776 | 8317 | 5298625 | -2638008.645 | 2681 | 5312848 |
| | 154 | -269601.7756 | 822 | 548603 | -274501.0773 | 290 | 552318 |
| | 15 | -19264.4287 | 71 | 39154 | -19465.3026 | 50 | 39371 |
| Bovidae | 15,486 | -14276063.64 | 60132 | 29516003 | -14675053.78 | 16919 | 29621308 |
| | 1,548 | -1419112.739 | 5975 | 2920210 | -1458435.435 | 1711 | 2940348 |
| | 154 | -144047.1052 | 562 | 294516 | -147514.636 | 185 | 297143 |
| | 15 | -8365.8073 | 64 | 17284 | -8579.0994 | 35 | 17460 |
| Carnivora | 15,485 | -20189220.63 | 73591 | 41558056 | -20634927.77 | 22881 | 41636623 |
| | 1,548 | -2042901.629 | 7358 | 4186822 | -2088967.935 | 2306 | 4209595 |
| | 154 | -223544.4981 | 766 | 455903 | -228291.2125 | 252 | 459482 |
| | 15 | -20813.3401 | 72 | 42298 | -21110.0594 | 42 | 42612 |
| Myomorpha | 15,486 | -25576581.35 | 82612 | 52477370 | -25988907.4 | 26527 | 52403022 |
| | 1,548 | -2479841.109 | 8281 | 5073203 | -2521869.523 | 2692 | 5080643 |
| | 154 | -232786.927 | 841 | 475139 | -236472.9178 | 284 | 476176 |
| | 15 | -25439.8935 | 97 | 51768 | -25802.2226 | 49 | 52053 |
| Primates | 15,486 | -25127036.02 | 79361 | 51526169 | -25556510.21 | 25658 | 51524299 |
| | 1,548 | -2531565.112 | 8046 | 5173550 | -2575829.296 | 2631 | 5187765 |
| | 154 | -256432.8736 | 841 | 522495 | -261055.6783 | 306 | 525615 |
| | 15 | -17934.338 | 109 | 36852 | -18205.7104 | 65 | 36998 |

### FIGURES

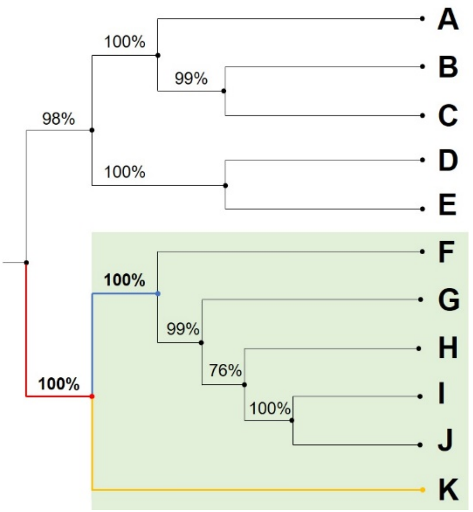


FIGURE A.1. ML tree with the bootstrap support values for each branch. The only clade that contains at least five taxa and has 100% bootstrap support at the branch leading to that clade and at the first direct descendants in the clade is the green tree (F-K). The red branch is the branch leading to the clade, the blue and the yellow branches are the descendant branches. Note, since the yellow branch is a tip, the bootstrap support is 100%.

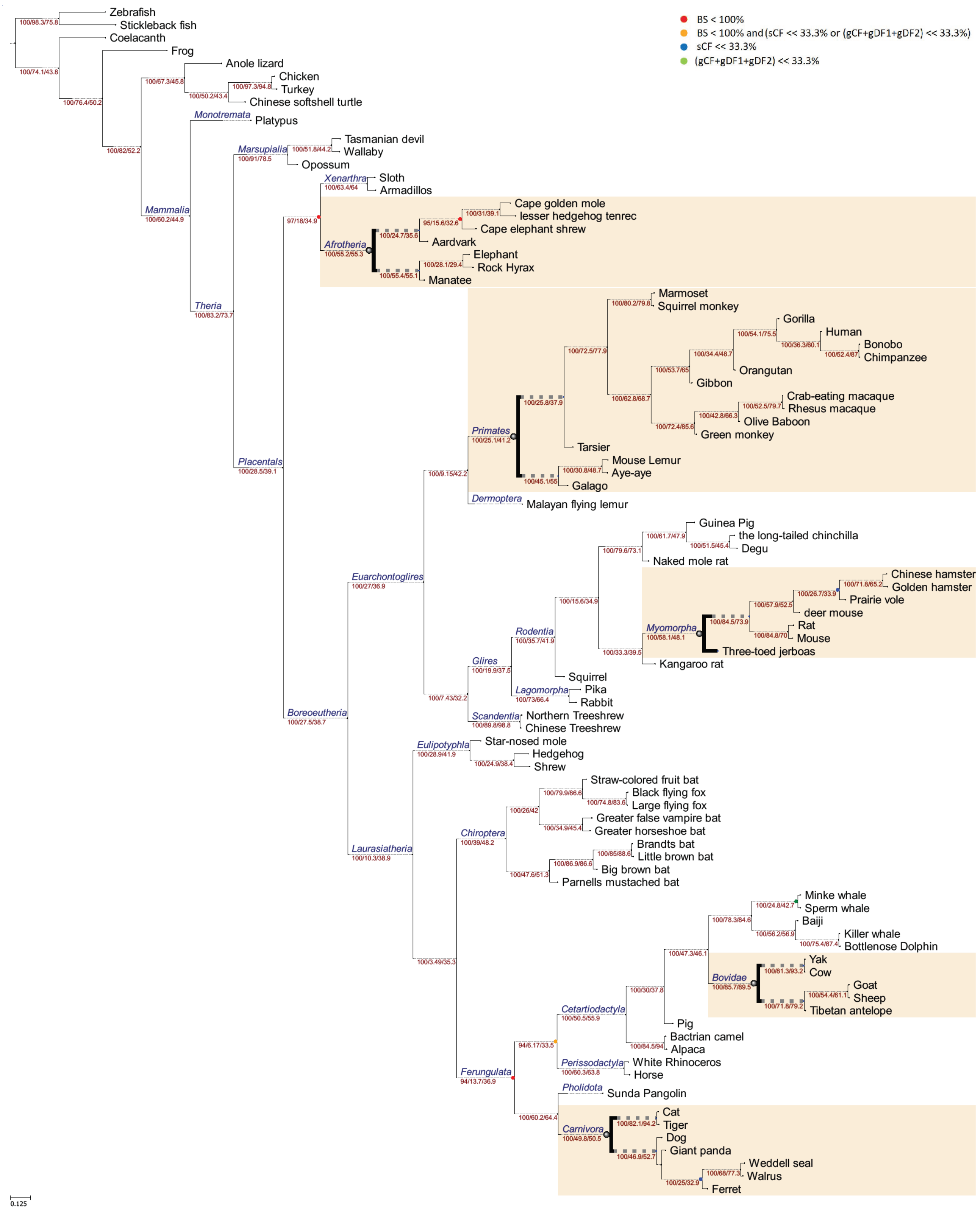


FIGURE A.2. The ML tree inferred from the whole concatenated AA alignment from (Wu, et al. 2018) and rooted on non-mammalian outgroup taxa. Bold branches present the well-defined clades we use in this study.

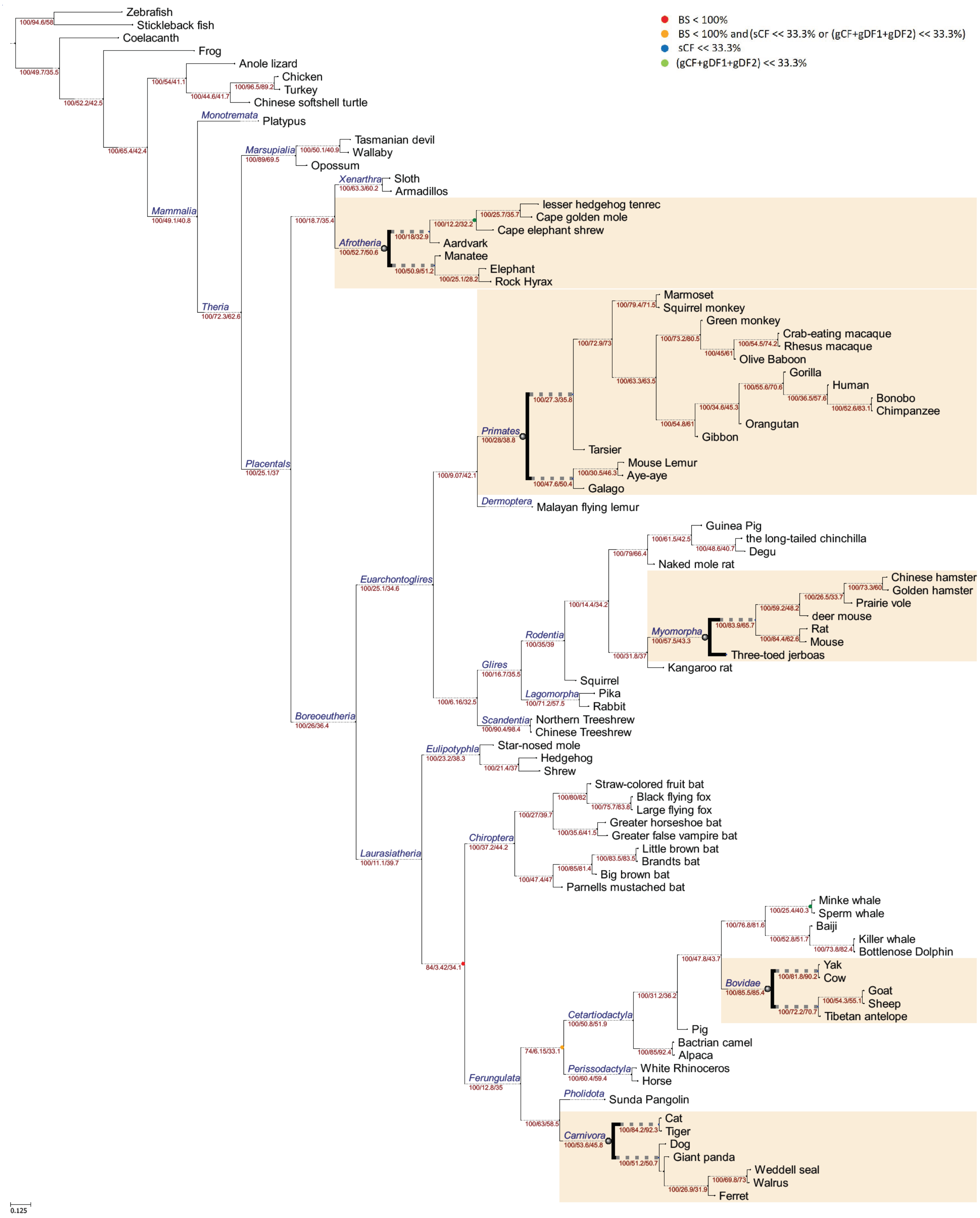


FIGURE A.3. The ML tree inferred from the whole concatenated DNA alignment from (Wu, et al. 2018) and rooted on non-mammalian outgroup taxa.

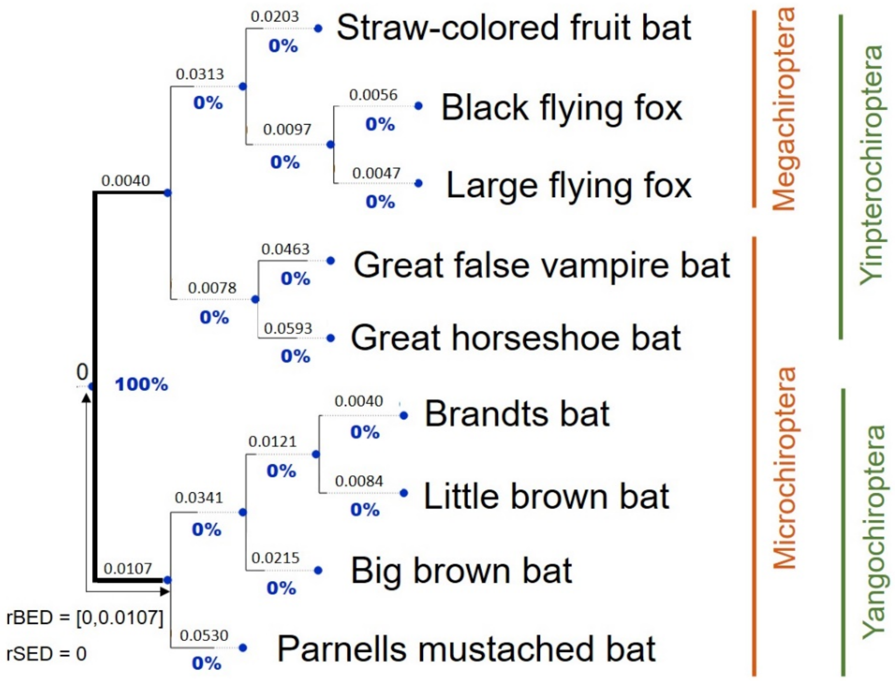


FIGURE A.4. The ML rooted tree of as inferred from the whole Chiroptera nucleotide dataset. Bold branches are branches in the AU confidence set. Blue values under each branch are the rootstrap support values.

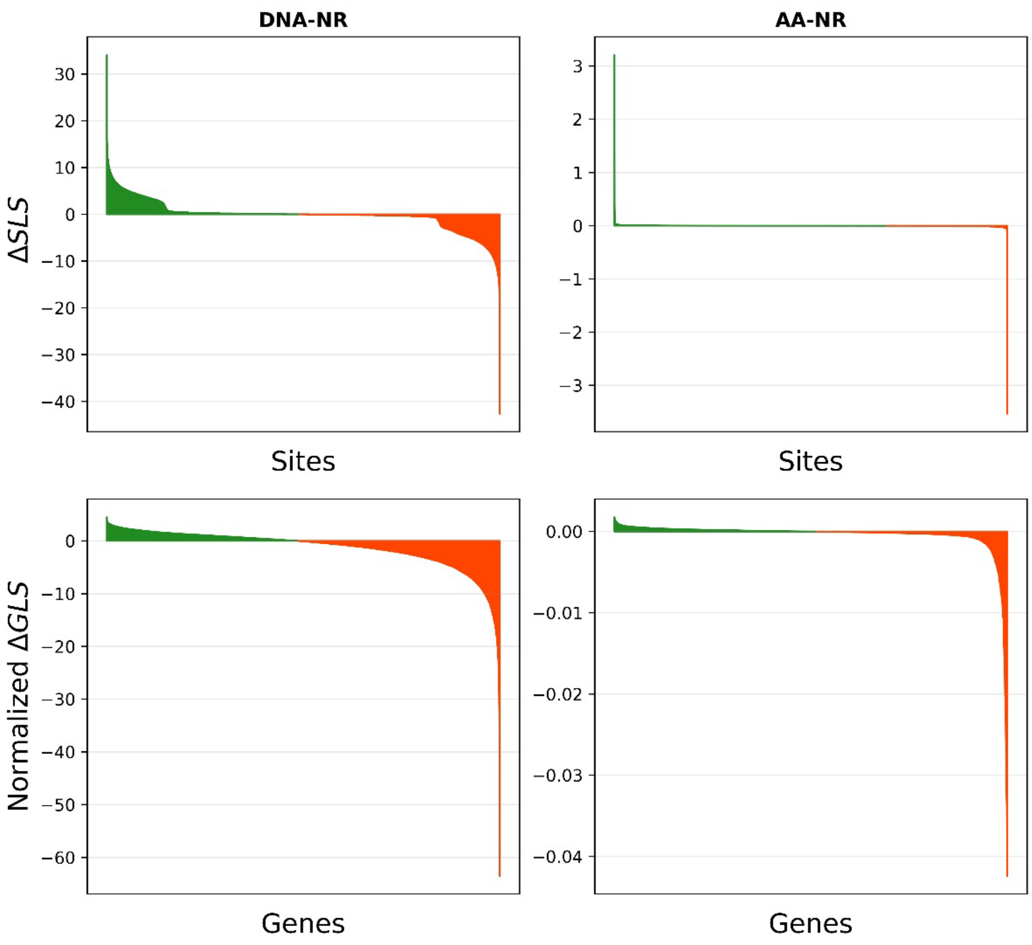


FIGURE A.5. The normalized difference in the gene-wise log-likelihood score (ΔGLS) and the difference in the site-wise log-likelihood score (ΔSLS) in the Chiroptera amino acid and nucleotide datasets. Positive values (green) present genes/sites that favour the Yinptero-Yango hypothesis and negative values present genes/sites that favour the Micro-Mega hypothesis.

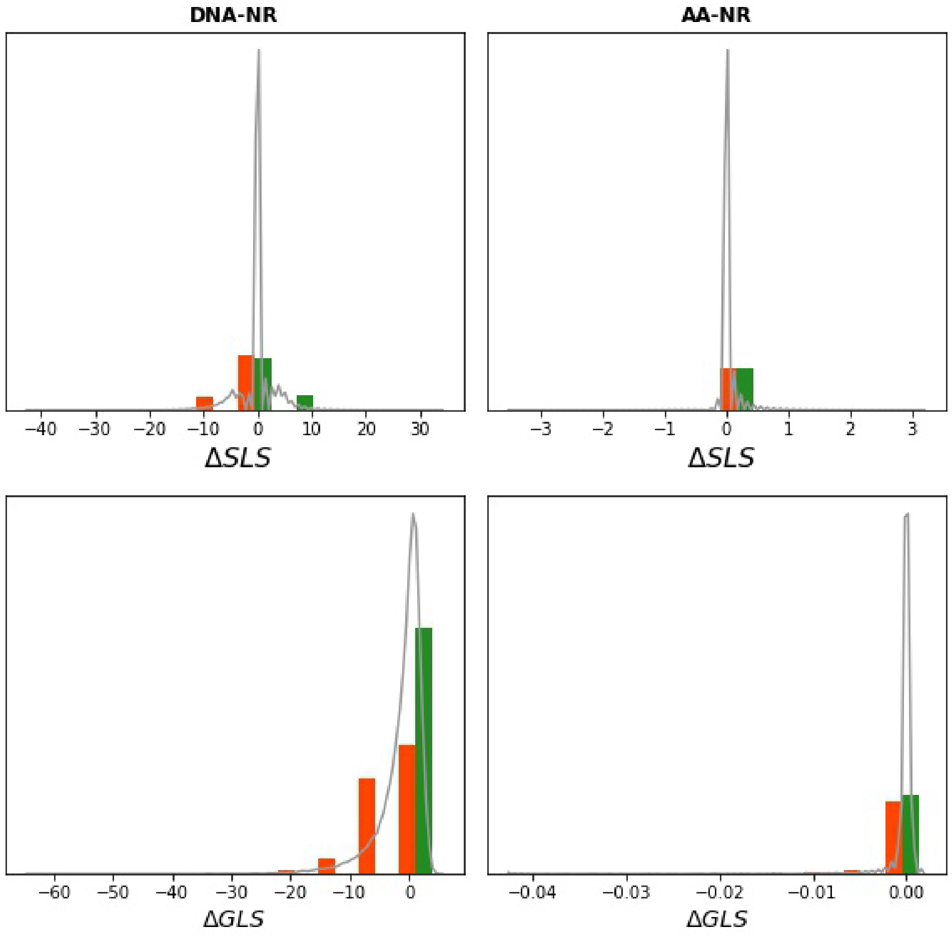


FIGURE A.6. The distribution of the normalized ΔGLS and ΔSLS in the Chiroptera amino acid and nucleotide datasets where green presents genes that support the Yinptero-Yango hypothesis and orange presents genes that support the Micro-Mega hypothesis.

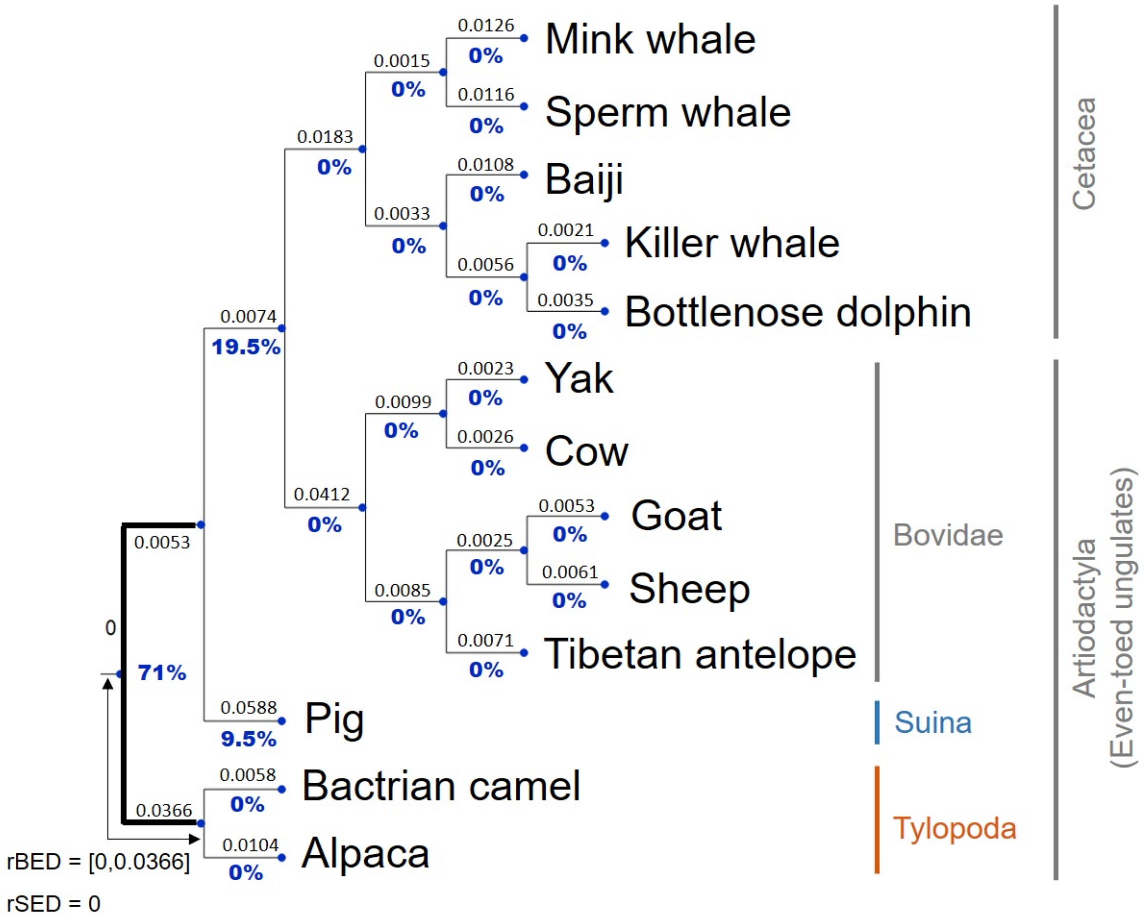


FIGURE A.7. The ML rooted tree of as inferred from the whole Cetartiodactyla nucleotide dataset. Bold branches are branches in the AU confidence set. Blue values under each branch are the rootstrap support values.

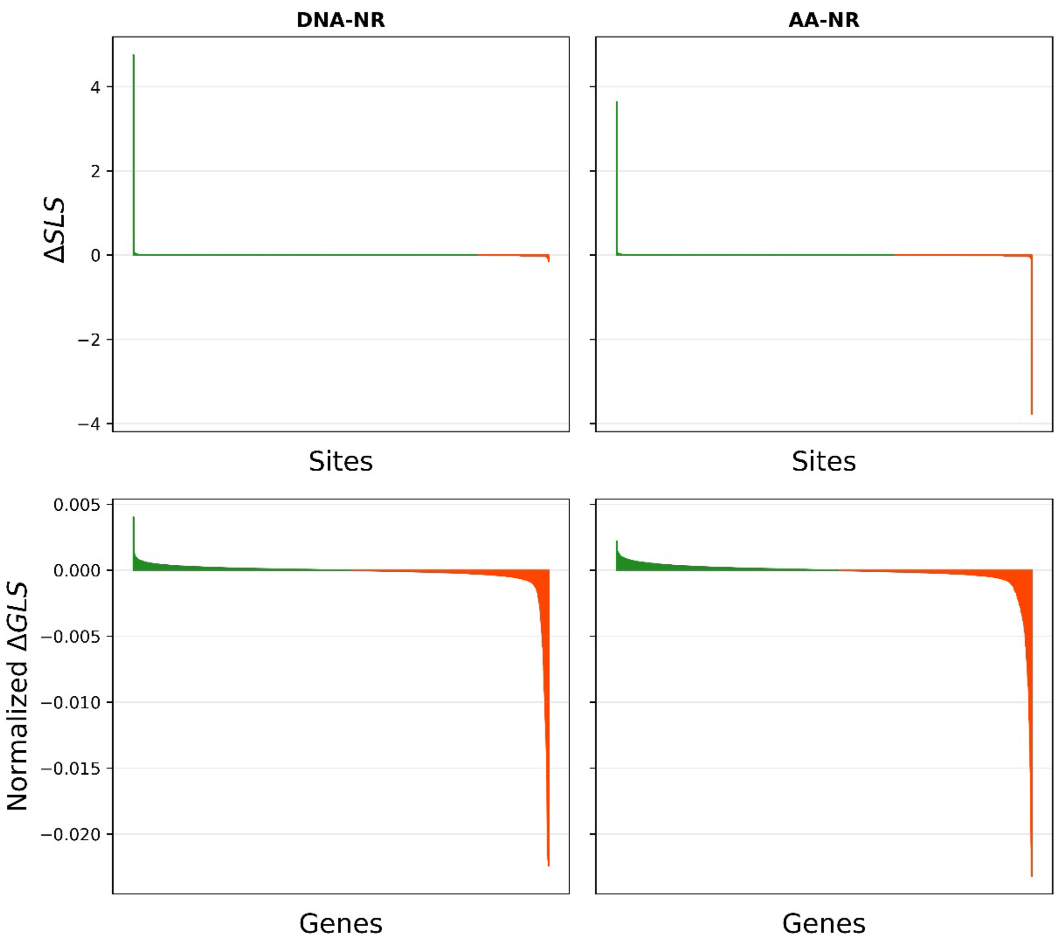


FIGURE A.8. The normalized difference in the gene-wise log-likelihood score (ΔGLS) and the difference in the site-wise log-likelihood score (ΔSLS) between the Tylopoda+Suina hypothesis and the Tylopoda hypothesis. Positive values (green) present genes/sites that favour the Tylopoda+Suina hypothesis and negative values (orange) present genes/sites that favour the Tylopoda hypothesis.

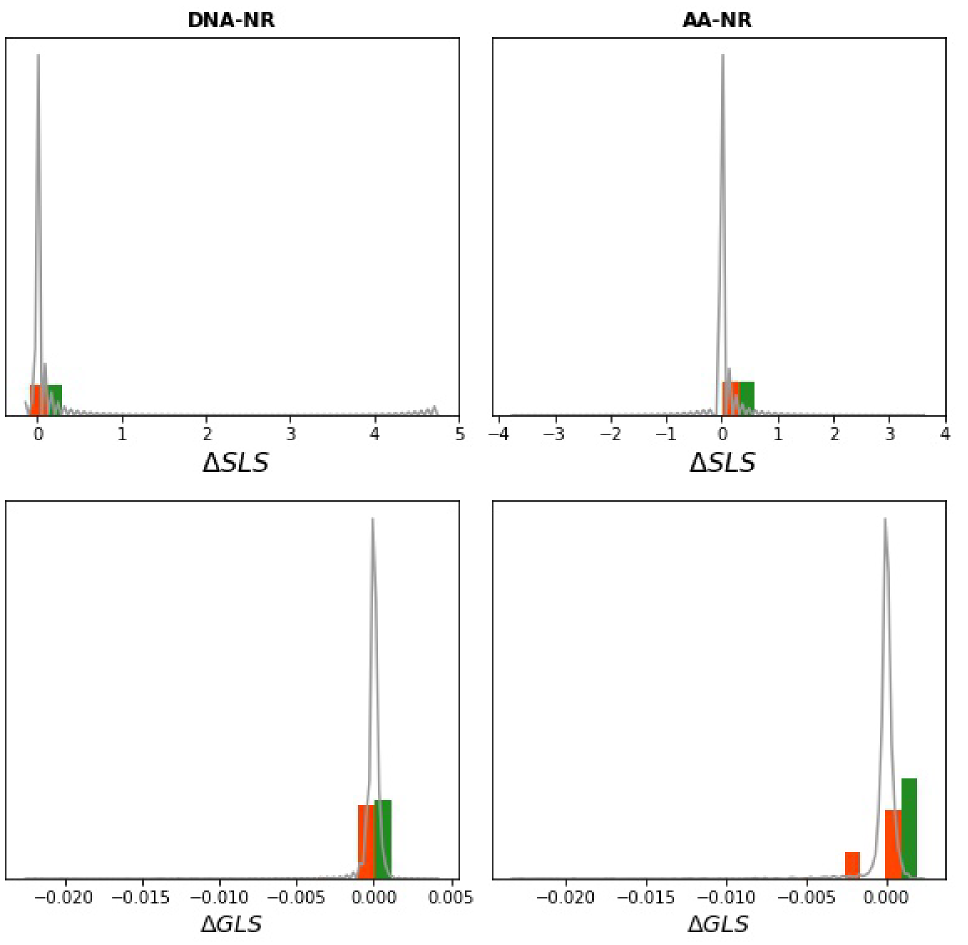


FIGURE A.9. The distribution of the normalized ΔGLS and ΔSLS in in the Cetartiodactyla amino acid and nucleotide dataset where green presents genes that support the Tylopoda+Suina hypothesis and orange presents genes that support the Tylopoda hypothesis.

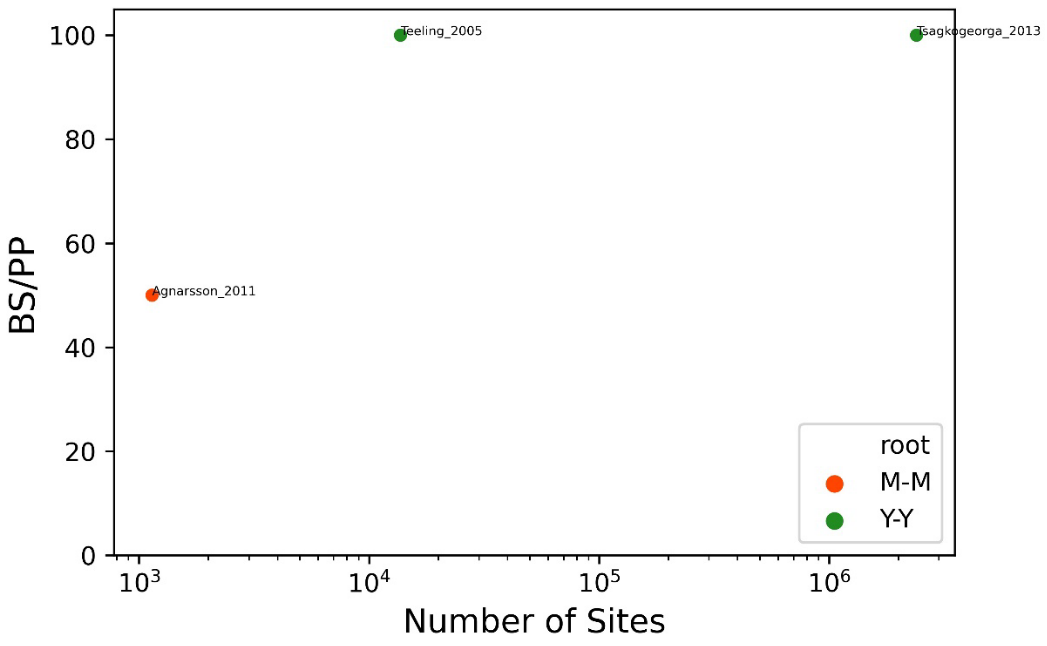


FIGURE A.10. The support value (bootstrap or posterior probability) for a root placement against the number of nucleotide sites used in each study. Orange dot is for Micro-Mega hypothesis (Agnarsson, et al. 2010) and green dots are for the Yinpterochiroptera-Yangochiroptera hypothesis (Teeling, et al. 2005; Tsagkogeorga, et al. 2013).

### REFERENCES

Agnarsson I, Kuntner M, May-Collado LJ. 2010. Dogs, cats, and kin: a molecular species-level phylogeny of Carnivora. Mol Phylogenet Evol 54:726-745.

Asher RJ, Bennett N, Lehmann T. 2009. The new framework for understanding placental mammal evolution. Bioessays 31:853-864.

Benoit J, Crumpton N, Merigeaud S, Tabuce R. 2013. A memory already like an elephant's? The advanced brain morphology of the last common ancestor of Afrotheria (Mammalia). Brain Behav. Evol. 81:154-169.

Bibi F. 2013. A multi-calibrated mitochondrial phylogeny of extant Bovidae (Artiodactyla, Ruminantia) and the importance of the fossil record to systematics. BMC Evol. Biol. 13:166.

Bibi F, Bukhsianidze M, Gentry AW, Geraads D, Kostopoulos DS, Vrba ES. 2009. The Fossil Record and Evolution of Bovidae: State of the Field. Palaeontologia Electronica 12:1-11.

Blanga-Kanfi S, Miranda H, Penn O, Pupko T, DeBry RW, Huchon D. 2009. Rodent phylogeny revised: analysis of six nuclear genes from all major rodent clades. BMC Evol. Biol. 9:71.

Chatterjee HJ, Ho SY, Barnes I, Groves C. 2009. Estimating the phylogeny and divergence times of primates using a supermatrix approach. BMC Evol. Biol. 9:259.

Chen L, Qiu Q, Jiang Y, Wang K, Lin Z, Li Z, Bibi F, Yang Y, Wang J, Nie W, et al. 2019. Large-scale ruminant genome sequencing provides insights into their evolution and distinct traits. Science 364:eaav6202.

Churakov G, Sadasivuni MK, Rosenbloom KR, Huchon D, Brosius J, Schmitz J. 2010. Rodent evolution: back to the root. Mol. Biol. Evol. 27:1315-1326.

Diogo R, Wood B. 2011. Soft-tissue anatomy of the primates: phylogenetic analyses based on the muscles of the head, neck, pectoral region and upper limb, with notes on the evolution of these muscles. J. Anat. 219:273-359.

dos Reis M, Inoue J, Hasegawa M, Asher RJ, Donoghue PC, Yang Z. 2012. Phylogenomic datasets provide both precision and accuracy in estimating the timescale of placental mammal phylogeny. Proc Biol Sci 279:3491-3500.

Eizirik E, Murphy WJ, Koepfli KP, Johnson WE, Dragoo JW, Wayne RK, O'Brien SJ. 2010. Pattern and timing of diversification of the mammalian order Carnivora inferred from multiple nuclear gene sequences. Mol Phylogenet Evol 56:49-63.

Elliot MG, Crespi BJ. 2009. Phylogenetic evidence for early hemochorial placentation in eutheria. Placenta 30:949-967.

Fabre PH, Hautier L, Dimitrov D, Douzery EJ. 2012. A glimpse on the pattern of rodent diversification: a phylogenetic approach. BMC Evol. Biol. 12:88.

Fabre PH, Rodrigues A, Douzery EJ. 2009. Patterns of macroevolution among Primates inferred from a supermatrix of mitochondrial and nuclear DNA. Mol Phylogenet Evol 53:808-825.

Finarelli JA, Flynn JJ. 2009. Brain-size evolution and sociality in Carnivora. Proc Natl Acad Sci U S A 106:9345-9349.

Finstermeier K, Zinner D, Brameier M, Meyer M, Kreuz E, Hofreiter M, Roos C. 2013. A mitogenomic phylogeny of living primates. PLoS One 8:e69504.

Foley NM, Springer MS, Teeling EC. 2016. Mammal madness: is the mammal tree of life not yet resolved? Philos Trans R Soc Lond B Biol Sci 371:20150140.

Gheerbrant E, Amaghzaz M, Bouya B, Goussard F, Letenneur C. 2014. Ocepeia (Middle Paleocene of Morocco): the oldest skull of an afrotherian mammal. PLoS One 9:e89739.

Halliday TJ, Upchurch P, Goswami A. 2015. Resolving the relationships of Paleocene placental mammals. Biol. Rev. Camb. Philos. Soc. 92:521-550.

Hao H, Liu S, Zhang X, Chen W, Song Z, Peng H, Liu Y, Yue B. 2011. Complete mitochondrial genome of a new vole Proedromys liangshanensis (Rodentia: Cricetidae) and phylogenetic analysis with related species: are there implications for the validity of the genus Proedromys? Mitochondrial DNA 22:28-34.

Hartig G, Churakov G, Warren WC, Brosius J, Makalowski W, Schmitz J. 2013. Retrophylogenomics place tarsiers on the evolutionary branch of anthropoids. Sci Rep 3:1756.

Hassanin A, Delsuc F, Ropiquet A, Hammer C, Jansen van Vuuren B, Matthee C, Ruiz-Garcia M, Catzeflis F, Areskoug V, Nguyen TT, et al. 2012. Pattern and timing of diversification of Cetartiodactyla (Mammalia, Laurasiatheria), as revealed by a comprehensive analysis of mitochondrial genomes. C. R. Biol. 335:32-50.

Hedrick BP, Dickson BV, Dumont ER, Pierce SE. 2020. The evolutionary diversity of locomotor innovation in rodents is not linked to proximal limb morphology. Sci Rep 10:717.

Herrera JP, Davalos LM. 2016. Phylogeny and Divergence Times of Lemurs Inferred with Recent and Ancient Fossils in the Tree. Syst. Biol. 65:772-791.

Hoang DT, Chernomor O, von Haeseler A, Minh BQ, Vinh LS. 2018. UFBoot2: Improving the Ultrafast Bootstrap Approximation. Mol. Biol. Evol. 35:518-522.

Horn S, Durka W, Wolf R, Ermala A, Stubbe A, Stubbe M, Hofreiter M. 2011. Mitochondrial genomes reveal slow rates of molecular evolution and the timing of speciation in beavers (Castor), one of the largest rodent species. PLoS One 6:e14622.

Jameson NM, Hou ZC, Sterner KN, Weckle A, Goodman M, Steiper ME, Wildman DE. 2011. Genomic data reject the hypothesis of a prosimian primate clade. J. Hum. Evol. 61:295-305.

Kari L, Hill KA, Sayem AS, Karamichalis R, Bryans N, Davis K, Dattani NS. 2015. Mapping the space of genomic signatures. PLoS One 10:e0119815.
Kishino H, Hasegawa M. 1989. Evaluation of the maximum likelihood estimate of the evolutionary tree topologies from DNA sequence data, and the branching order in hominoidea. J. Mol. Evol. 29:170-179.
Kishino H, Miyata T, Hasegawa M. (Kishino1990 co-authors). 1990. Maximum likelihood inference of protein phylogeny and the origin of chloroplasts. J. Mol. Evol. 31:151-160.
Kumar V, Hallstrom BM, Janke A. 2013. Coalescent-based genome analyses resolve the early branches of the euarchontoglires. PLoS One 8:e60019.
Kuntner M, May-Collado LJ, Agnarsson I. 2011. Phylogeny and conservation priorities of afrotherian mammals (Afrotheria, Mammalia). Zool. Scr. 40:1-15.
Lartillot N, Delsuc F. 2012. Joint reconstruction of divergence times and life-history evolution in placental mammals using a phylogenetic covariance model. Evolution 66:1773-1787.
Liu L, Zhang J, Rheindt FE, Lei F, Qu Y, Wang Y, Zhang Y, Sullivan C, Nie W, Wang J, et al. 2017. Genomic evidence reveals a radiation of placental mammals uninterrupted by the KPg boundary. Proc Natl Acad Sci U S A 114:E7282-E7290.
Machado FA, Zahn TMG, Marroig G. 2018. Evolution of morphological integration in the skull of Carnivora (Mammalia): Changes in Canidae lead to increased evolutionary potential of facial traits. Evolution 72:1399-1419.
Matsui A, Rakotondraparany F, Munechika I, Hasegawa M, Horai S. 2009. Molecular phylogeny and evolution of prosimians based on complete sequences of mitochondrial DNAs. Gene 441:53-66.
Meredith RW, Janecka JE, Gatesy J, Ryder OA, Fisher CA, Teeling EC, Goodbla A, Eizirik E, Simao TL, Stadler T, et al. 2011. Impacts of the Cretaceous Terrestrial Revolution and KPg extinction on mammal diversification. Science 334:521-524.
Minh BQ, Hahn M, Lanfear R. 2018. New methods to calculate concordance factors for phylogenomic datasets. bioRxiv:487801.

Monson TA, Hlusko LJ. 2018. Breaking the rules: Phylogeny, not life history, explains dental eruption sequence in primates. Am. J. Phys. Anthropol. 167:217-233.

Nyakatura K, Bininda-Emonds OR. 2012. Updating the evolutionary history of Carnivora (Mammalia): a new species-level supertree complete with divergence time estimates. BMC Biol. 10:12.

Panciroli E, Janis C, Stockdale M, Martin-Serra A. 2017. Correlates between calcaneal morphology and locomotion in extant and extinct carnivorous mammals. J. Morphol. 278:1333-1353.

Pattinson DJ, Thompson RS, Piotrowski AK, Asher RJ. 2015. Phylogeny, paleontology, and primates: do incomplete fossils bias the tree of life? Syst. Biol. 64:169-186.

Perelman P, Johnson WE, Roos C, Seuanez HN, Horvath JE, Moreira MA, Kessing B, Pontius J, Roelke M, Rumpler Y, et al. 2011. A molecular phylogeny of living primates. PLoS Genet. 7:e1001342.

Phillips MJ, Penny D. 2010. Mammalian Phylogeny. In. Encyclopedia of Life Sciences.

Polly PD, Fuentes-Gonzalez J, Lawing AM, Bormet AK, Dundas RG. 2017. Clade sorting has a greater effect than local adaptation on ecometric patterns in Carnivora. Evol. Ecol. Res. 18:61-95.

Poulakakis N, Stamatakis A. 2010. Recapitulating the evolution of Afrotheria: 57 genes and rare genomic changes (RGCs) consolidate their history. Syst. Biodivers. 8:395-408.

Pozzi L, Hodgson JA, Burrell AS, Sterner KN, Raaum RL, Disotell TR. 2014. Primate phylogenetic relationships and divergence dates inferred from complete mitochondrial genomes. Mol Phylogenet Evol 75:165-183.

Puttick MN, Thomas GH. 2015. Fossils and living taxa agree on patterns of body mass evolution: a case study with Afrotheria. Proc Biol Sci 282:20152023.

Reis MD, Gunnell GF, Barba-Montoya J, Wilkins A, Yang Z, Yoder AD. 2018. Using Phylogenomic Data to Explore the Effects of Relaxed Clocks and Calibration Strategies on Divergence Time Estimation: Primates as a Test Case. Syst. Biol. 67:594-615.

Romiguier J, Ranwez V, Douzery EJ, Galtier N. 2010. Contrasting GC-content dynamics across 33 mammalian genomes: relationship with life-history traits and chromosome sizes. Genome Res. 20:1001-1009.

Schenk JJ, Rowe KC, Steppan SJ. 2013. Ecological opportunity and incumbency in the diversification of repeated continental colonizations by muroid rodents. Syst. Biol. 62:837-864.

Shimodaira H. 2002. An approximately unbiased test of phylogenetic tree selection. Syst. Biol. 51:492-508.

Shimodaira H, Hasegawa M. 1999. Multiple comparisons of log-likelihoods with applications to phylogenetic inference. Mol. Biol. Evol. 16:1114-1116.

Springer MS, Meredith RW, Gatesy J, Emerling CA, Park J, Rabosky DL, Stadler T, Steiner C, Ryder OA, Janecka JE, et al. 2012. Macroevolutionary dynamics and historical biogeography of primate diversification inferred from a species supermatrix. PLoS One 7:e49521.

Stankowich T, Caro T, Cox M. 2011. Bold coloration and the evolution of aposematism in terrestrial carnivores. Evolution 65:3090-3099.

Steiper ME, Seiffert ER. 2012. Evidence for a convergent slowdown in primate molecular rates and its implications for the timing of early primate evolution. Proc Natl Acad Sci U S A 109:6006-6011.

Svartman M, Stanyon R. 2012. The chromosomes of Afrotheria and their bearing on mammalian genome evolution. Cytogenet. Genome Res. 137:144-153.

Swanson MT, Oliveros CH, Esselstyn JA. 2019. A phylogenomic rodent tree reveals the repeated evolution of masseter architectures. Proc Biol Sci 286:20190672.

Tavares WC, Seuanez HN. 2018. Changes in selection intensity on the mitogenome of subterranean and fossorial rodents respective to aboveground species. Mamm. Genome 29:353-363.

Teeling EC, Springer MS, Madsen O, Bates P, O'Brien S J, Murphy WJ. 2005. A molecular phylogeny for bats illuminates biogeography and the fossil record. Science 307:580-584.

Tomiya S, Tseng ZJ. 2016. Whence the beardogs? Reappraisal of the Middle to Late Eocene 'Miacis' from Texas, USA, and the origin of Amphicyonidae (Mammalia, Carnivora). R Soc Open Sci 3:160518.

Tsagkogeorga G, Parker J, Stupka E, Cotton JA, Rossiter SJ. 2013. Phylogenomic analyses elucidate the evolutionary relationships of bats. Curr. Biol. 23:2262-2267.

Wu J, Hasegawa M, Zhong Y, Yonezawa T. 2014. Importance of synonymous substitutions under dense taxon sampling and appropriate modeling in reconstructing the mitogenomic tree of Eutheria. Genes Genet. Syst. 89:237-251.

Wu J, Yonezawa T, Kishino H. 2017. Rates of Molecular Evolution Suggest Natural History of Life History Traits and a Post-K-Pg Nocturnal Bottleneck of Placentals. Curr. Biol. 27:3025-3033 e3025.

Wu S, Edwards S, Liu L. 2018. Genome-scale DNA sequence data and the evolutionary history of placental mammals. Data Brief 18:1972-1975.

Wu S, Wu W, Zhang F, Ye J, Ni X, Sun J, Edwards SV, Meng J, Organ CL. 2012. Molecular and paleontological evidence for a post-Cretaceous origin of rodents. PLoS One 7:e46445.

Yang C, Xiang C, Qi W, Xia S, Tu F, Zhang X, Moermond T, Yue B. 2013. Phylogenetic analyses and improved resolution of the family Bovidae based on complete mitochondrial genomes. Biochem. Syst. Ecol. 48:136-143.

Yue H, Yan CC, Tu FY, Yang CZ, Ma WQ, Fan ZX, Song ZB, Owens J, Liu SY, Zhang XY. 2015. Two novel mitogenomes of Dipodidae species and phylogeny of Rodentia inferred from the complete mitogenomes. Biochem. Syst. Ecol. 60:123-130.

Zhang ML, Li ML, Ayoola AO, Murphy RW, Wu DD, Shao Y. 2019. Conserved sequences identify the closest living relatives of primates. Zool. Res. 40:532-540.
